# Pronounced sequence specificity of the TET enzyme catalytic domain guides its cellular function

**DOI:** 10.1101/2021.12.29.474486

**Authors:** Mirunalini Ravichandran, Dominik Rafalski, Oscar Ortega-Recalde, Claudia I. Davies, Cassandra R. Glanfield, Annika Kotter, Katarzyna Misztal, Andrew H. Wang, Marek Wojciechowski, Michał Rażew, Issam M. Mayyas, Olga Kardailsky, Uwe Schwarz, Krzysztof Zembrzycki, Ian M. Morison, Mark Helm, Dieter Weichenhan, Renata Z. Jurkowska, Felix Krueger, Christoph Plass, Martin Zacharias, Matthias Bochtler, Timothy A. Hore, Tomasz P. Jurkowski

## Abstract

TET (ten-eleven translocation) enzymes catalyze the oxidation of 5-methylcytosine bases in DNA, thus driving active and passive DNA demethylation. Here, we report that the catalytic cores of mammalian TET enzymes favor CpGs embedded within bHLH and bZIP transcription factor binding sites, with 250-fold preference *in vitro*. Crystal structures and molecular dynamics calculations show that sequence preference is caused by intra-substrate interactions and CpG flanking sequence indirectly affecting enzyme conformation. TET sequence preferences are physiologically relevant as they explain the rates of DNA demethylation in TET-rescue experiments in culture and *in vivo* within the zygote and germline. Most and least favorable TET motifs represent DNA sites that are bound by methylation-sensitive immediate-early transcription factors and OCT4, respectively, illuminating TET function in transcriptional responses and pluripotency support.

**One-Sentence Summary:** The catalytic domains of the enzymes that facilitate passive and drive active DNA demethylation have intrinsic sequence preferences that target DNA demethylation to bHLH and bZIP transcription factor binding sites.

## Main Text

DNA methylation in the form of 5-methylcytosine (5mC) is an epigenetic modification essential for mammalian development and cellular differentiation (*1*). TET enzymes catalyze the oxidation of 5mC bases in DNA to hydroxymethylcytosine (5hmC), formylcytosine (5fC), or carboxylcytosine (5caC) bases (*2, 3*). In doing so, TET proteins enhance the reprogramming of cultured cells to a pluripotent state (*4*–*6*), and allow the germline to achieve full developmental potency *in vivo* (*7*). TETs also play a role as tumor suppressors, judging from their frequent loss in acute myelogenic leukemia and other malignancies (*8*), further emphasizing their importance for modulating epigenetic regulation. TET oxidation favors both active, replication-uncoupled, and passive, replication-coupled DNA demethylation. Active DNA demethylation is primed by the oxidized 5-methylcytosine derivatives, which resemble damaged nucleobases (*9*). These are recognized and excised by the DNA repair machinery, particularly base excision repair, ultimately leading to the replacement of methylated 2’-deoxynucleotides by their unmodified congeners (*3*). Passive DNA demethylation is facilitated by the most abundant 5mC oxidation product, 5hmC, which prevents remethylation at the replisome of the daughter strand by the maintenance methyltransferase (*10*).

Although central to understanding TET function, the mechanism by which TET proteins are targeted to specific DNA sequences is not clear. CXXC or IDAX-mediated recruitment of TET proteins targets non-methylated CpGs, or more generally, regions of low cytosine methylation (*11*). While this may suggest TETs can function as epigenomic repair enzymes, it does not explain their strong demethylation capacity elsewhere. Post-translational modification is known to alter the binding of TET to DNA (*12*), and formation of TET complexes with transcription factors and other DNA binding proteins (*4, 13*) is thought to both alter TET targeting and support pluripotency. Histone marks and epigenomic chromatin states are good predictors for sites of TET action (*14*); however, this alone does not provide insight into how TET proteins select sites for catalysis.

Here, we show that the TET catalytic domain, previously considered solely a catalytic engine, significantly contributes to DNA target selection with a pronounced, up to 250-fold preference for some CpG sequence contexts over others. Moreover, both the most and least favorable motifs constitute methylation sensitive transcription factor binding sites and contribute a new understanding of TET enzyme function.

### Specificity of the TET catalytic domain *in vitro*

We initially discovered TET sequence preference in *in vitro* assays where recombinant catalytic domains of mouse TET1, 2, and 3 were incubated with libraries of DNA substrates where all cytosine positions were methylated (Figure 1A). In order to read out demethylation, a bisulfite assay that exploits the resistance of 5-methylcytosine, but not its oxidation products 5-formyl-cytosine and 5-carboxylcytosine, to bisulfite driven deamination, which manifests itself, after PCR amplification, as a C->T transition (Figures 1B, C and S1B). As expected, non-CG sequences were demethylated at a much slower rate than CG sites (Figures 1B, 2A and 2B), with decreasing activity towards CG>>CC>CA>CT (for TET3 and TET1). Among the CG-containing sequences, we uncovered a 250-fold dynamic range between the most rapidly and the most slowly demethylated sites. The top-ranked demethylating sequence was the CACGTG hexamer (*in vitro* demethylation velocity 0.058±0.008 fraction converted per minute). Importantly, this sequence represents the canonical E-box motif, a well-known recognition site for many helix-loop-helix (bHLH) and basic zipper leucine domain (bZIP) transcription factors, the most iconic of which is the c-MYC oncogene. Many of the other rapidly demethylated sequences had an adenine upstream (−1 position) and a thymine downstream (+1 position) of the CG, with 7 out of the top 13 (53.8%) fastest demethylating sites possessed this motif. At the other end of the spectrum, the most slowly demethylated sequence was GGCGGG (0.00023±0.00009 fraction converted per minute).

**Figure 1.**
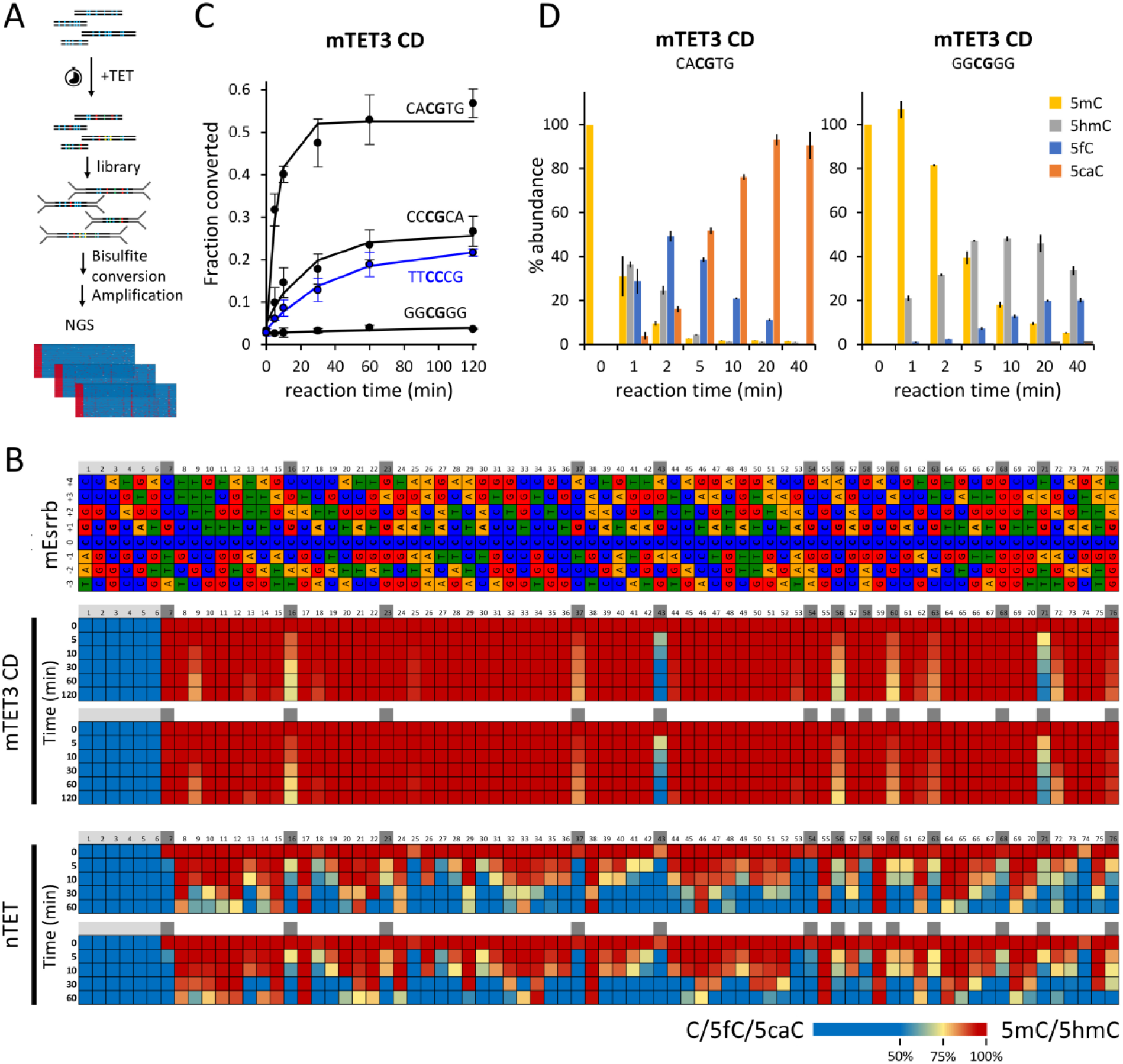
*In vitro* flanking sequence preference of TET enzymes. **(A)** Outline of the *in vitro* demethylation kinetics setup. 5mC-modified dsDNA substrates were incubated with recombinant TET catalytic domains for various lengths of time, products were purified and ligated to Illumina adapters, bisulfite converted and sequenced. **(B)** TET activity profile on fully 5mC-modified mouse Esrrb (mEsrrb) promoter fragment. The seven Cs on the 5’ end of the DNA substrates are devoid of modification, as these were part of the unmethylated primers used for substrate generation (colored with light grey). The rows represent the timepoints of reaction progression, the columns represent each potential 5mC site that could be modified. The color of the boxes denotes the methylation level of the site, according to the legend on the bottom of the panel. For convenience, CG sites are marked with grey boxes on top of the separate substrate panels. **(C)** Comparison of mTET3 CD reaction kinetics on CACGTG (fastest), CCCGCA (middle), GGCGGG (slowest) substrate identified in the screen. **(D)** Comparison of *in vitro* reaction kinetics of mTET3 oxidation of synthetic CACGTG (fastest) and GGCGGG (slowest) substrates measured using LCMS. For (C) and (D) error bars denote SD from two biological replicates.

Interestingly, demethylation of the least favorable CG sequences was even slower (23-fold slower) than for the most favorable non-CG substrates (GCCCTT; 0.0055± 0.00086 fraction converted per minute), suggesting that the flanking sequences strongly impact TET activity. We observed a similar sequence preference for mouse TET1 (Figures 2 and S1B) despite a lower overall activity observed in the assays.

**Figure 2.**
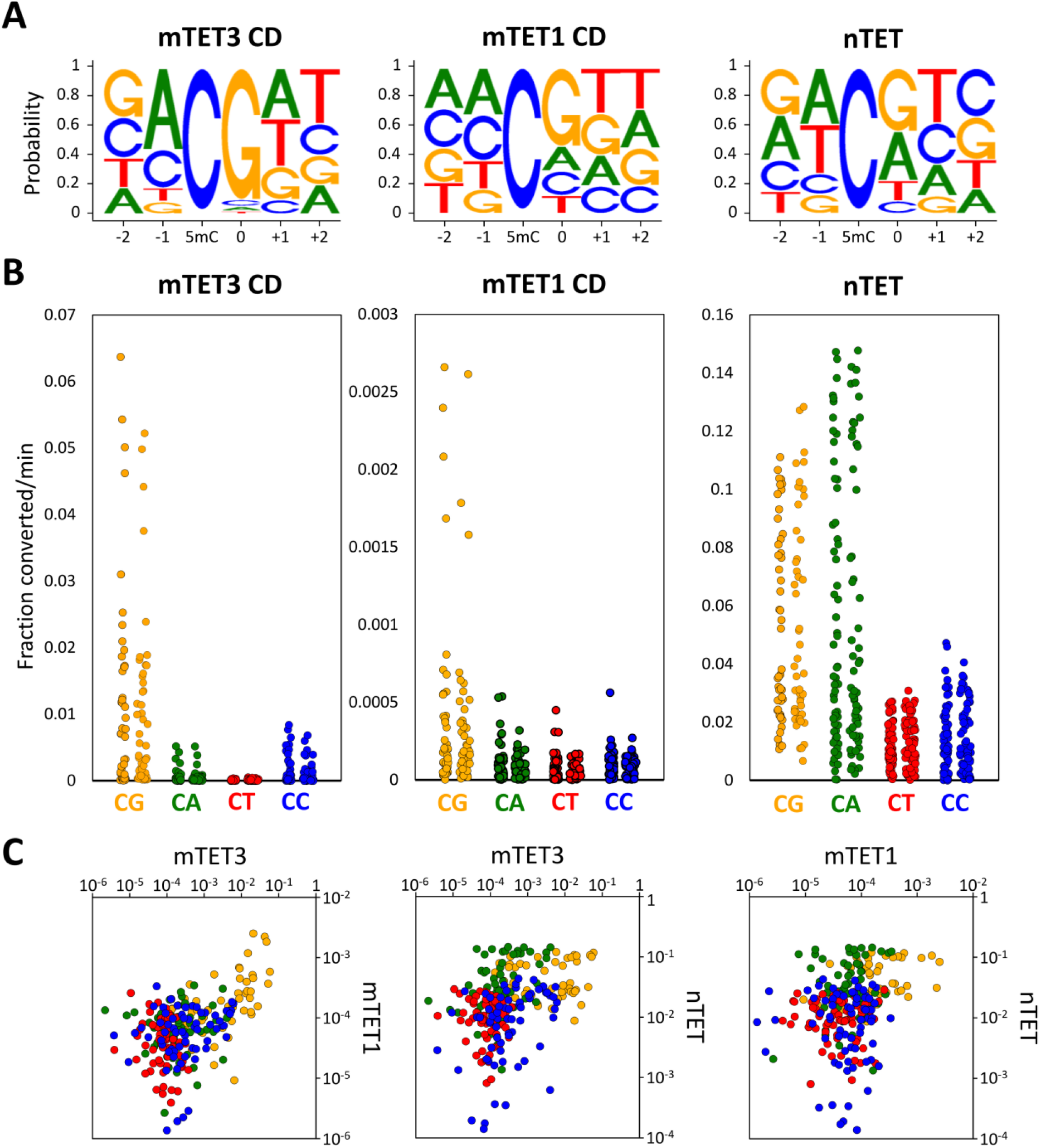
*In vitro* specificity profiles of TET enzymes. **(A)** Sequence logos of the TET enzymes based on the obtained *in vitro* reaction kinetics. **(B)** Dot plot of catalytic activities observed on 5mCs embedded in CG and non-CG contexts in all 4 tested *in vitro* substrates (mEsrrb, CG-rich, Tcl1 and Nanog). Each dot represents a 5mC site in either CG, CA, CT or CC context also differing in the flanking sequences beyond the central dinucleotide. **(C)** Pairwise comparison of activity profiles of mTET1 CD, mTET3 CD and nTET. Orange dots represent – CG, green – CA, red – CT and blue – CC sites. X and Y axis represent fraction converted 5mC per minute.

To independently corroborate these findings using an alternative *in vitro* assay, we compared the conversion of 5mCG in the E-box sequence (CACGTG) by TET3 to the most slowly demethylating motif identified in the screen (GGCGGG) using quantitative liquid-chromatography-coupled mass spectrometry (LC/MS) (Figure 1D). This analysis confirmed rapid transition of the central methylated cytosine within the E-box sequence to 5hmC (4.5%), 5fC (38.6%) and 5caC (51.8%) by 5 minutes into the reaction, whereas in the same time, the GGCGGG sequence had the central 5mC only oxidized to 5hmC (44%), 5fC (15.6%), 5caC (1.5%).

Furthermore, we compared the *in vitro* demethylation activity of mammalian TETs to that of the recombinant nTET enzyme from amoeba *Naegleria gruberi*. nTET is only remotely related to mammalian enzymes. It shares a similar overall structure yet is missing the Cys-rich region and binds DNA somewhat differently than mammalian TETs. nTET was shown to efficiently convert 5mC all the way to 5caC *in vitro* (*15*). We also observed a very robust catalytic activity of nTET on all the provided substrates and found that when compared to mammalian enzymes, nTET was able to oxidize 5mC in broader sequence context than mammalian TETs (Figures 1B and 2), preferentially oxidizing CA and CG sites over CC and CT sites, indicating that the sequence specificity profile obtained for TET1 and TET3 are characteristic for the mammalian enzymes.

To analyze sequence preferences quantitatively, we built predictive models for the demethylation rates, assuming an independent site model (i.e. overall preferences are products of individual site preferences). According to this model, logarithms of catalytic rates should be amenable to linear regression. This was indeed the case, with good correlation coefficients, indicating that preferences in the flanking regions of the central CG dinucleotide were not strongly interdependent (Figure S2). Overall, our *in vitro* results show that the activity of the mammalian TETs, and to a lesser extent also the nTET activity depends on sequence context flanking the target 5mC. The bases outside the central CG have a strong influence on the rates of catalysis. Interestingly, the TET1 and TET3 sequence preferences are similar, suggesting that the TET sequence preferences may be shared among paralogues (Figures 2 and S1B).

### Structural basis of TET sequence preference

To understand the structural basis for TET sequence preferences and their conservation among TET paralogues, we aimed to crystallize vertebrate TET protein complexes with the most and least favorable substrates. Among the vertebrate TET paralogues, only human TET2 could be crystallized in our hands. We used the truncated version of the protein (residues 1129–1936) with a 15-residue GS-linker replacing the internal disordered region (residues 1481-1843) similar to that used in the original report on the TET2 structure (*16*). We grew crystals in the previously published C222(1) crystal form, with oligoduplexes containing the most and least favorable sequences CA5mCGTG and GG5mCGCC, respectively. For crystallization, we used the enzyme with the native Fe^2+^ or Mn^2+^ in the active site and replaced the co-substrate 2-oxoglutarate with oxalylglycine, which does not support the reaction.

Overall, the structures with the two substrates are very similar to each other and resembled the previously published human TET2 co-structure (PDB accession: 4NM6) (*17*) (Figure S3 and 3B). Recognition of both the 5’- and 3’-flanking sequences is indirect, not due to direct discriminating amino acid contacts.

The TET sequence preference on the 5’-side of a CpG is due to interactions in the non-substrate strand. For the favorable substrate, the estranged guanine (that was originally base paired with the substrate 5mC) has its glycosidic bond to the 2’-deoxyribose in favorable anti-orientation and can donate a hydrogen bond from its exocyclic amino group to the T in −1 position in the bottom strand. By contrast, for the non-favorable substrate, the C in the bottom strand is in steric conflict with the estranged guanine in anti-orientation and therefore drives this nucleobase into the unfavorable *syn* conformation. A similar bottom strand steric conflict is also expected for G, but not A in the −1 position of the bottom strand. Therefore, the intra-strand interactions favor A or C (often abbreviated as M for a base with an exocyclic a<u>m</u>ino group) in the top strand in −1 position (Figure 3C).

**Figure 3.**
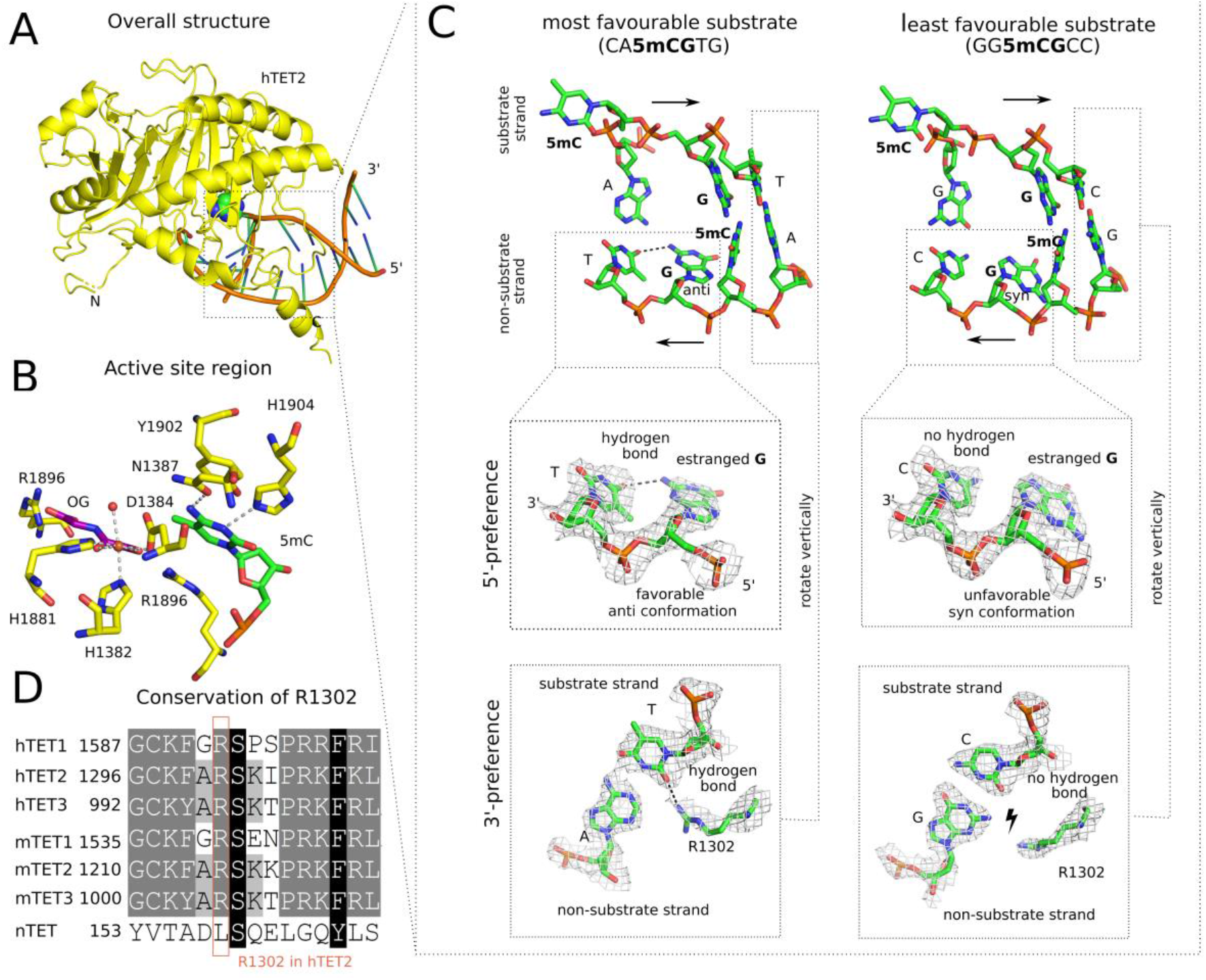
Structural basis for TET sequence specificity. **(A)** Structure of the core region of hTET2 (residues 1129–1936, with a 15-residue GS-linker replacing disordered residues 1481-1843) with the most favorable substrate. Protein is shown in yellow in ribbon representation, and DNA in schematic representation (brown backbone, green/blue nucleobases). The substrate 5mC base is highlighted in all-atom representation. The structure with the least favorable substrate is indistinguishable at this level of detail, except at the very N-terminus, which is very uncertain due to high B-factors (Figure S3). **(B)** Active site region with key active site residues (yellow), Fe^2+^ (brown), the co-substrate analog N-oxalylglycine (purple), and the substrate 5-methyl-2’-deoxycytidine monophosphate (green). At the level of resolution of the crystal structures, the active site regions are indistinguishable for the complexes with the most and least favorable substrates. **(C)** Conformation of the central four 2’-deoxynucleotides of substrate and non-substrate strands. In the magnified regions, composite omit densities contoured at 1 σ are shown. **(D)** Conservation of the arginine residue (R1302 in hTET2) responsible for 3’-substrate preferences in the TET paralogues.

The TET sequence preference on the 3’-side of the CpG is due to interactions with a conserved arginine residue (Arg1302 in TET2, Figure 3D). For the favorable A or T (often abbreviated as W for a base taking part in a <u>w</u>eak base pair), the arginine adopts an “in” conformation that enables formation of a hydrogen bond with a universal acceptor site in the minor groove of the DNA. By contrast, for the unfavorable C or G, the arginine is pushed into an “out” conformation by the presence of the 2-amino-group of guanine in the central minor groove, which abolishes the favorable interaction (Figure 3C).

Together, the biochemical and structural data suggest that TET sequence specificity is best described as MCGW. This conclusion was independently confirmed by molecular dynamics simulations, which generated very similar results to the crystallographic analysis, except for the transition to the disfavoured syn-conformation of the glycosidic bond for the unfavorable substrate that was not seen in the simulations (Figure S5).

### TET sequence specificity in culture and *in vivo*

To test the contribution of this inherent flanking sequence specificity of the TETs on the genomic demethylation pattern, we expressed a mouse TET3 catalytic domain transgene in mouse embryonic stem cells (mESCs) using an inducible piggyBAC construct (Figures 4A and S6A). To ensure there were no confounding interactions with endogenous TET proteins, the ESC line background used was a triple genetic knockout for TET1-3 (TET-TKO) (*18*). Methylation loss over 72 h was assessed by low-coverage whole-genome BS-seq in triplicate at 6 h time intervals following doxycycline (dox) induction (Figure 4A). We found that global CG methylation was reduced by 12.8 percentage points (pp) at 30 h post dox treatment and then slowly regained 3.0% by 72 h (red line, Figure 4B). In contrast, global methylation did not change in control cells over the same period (Figure S6B).

**Figure 4.**
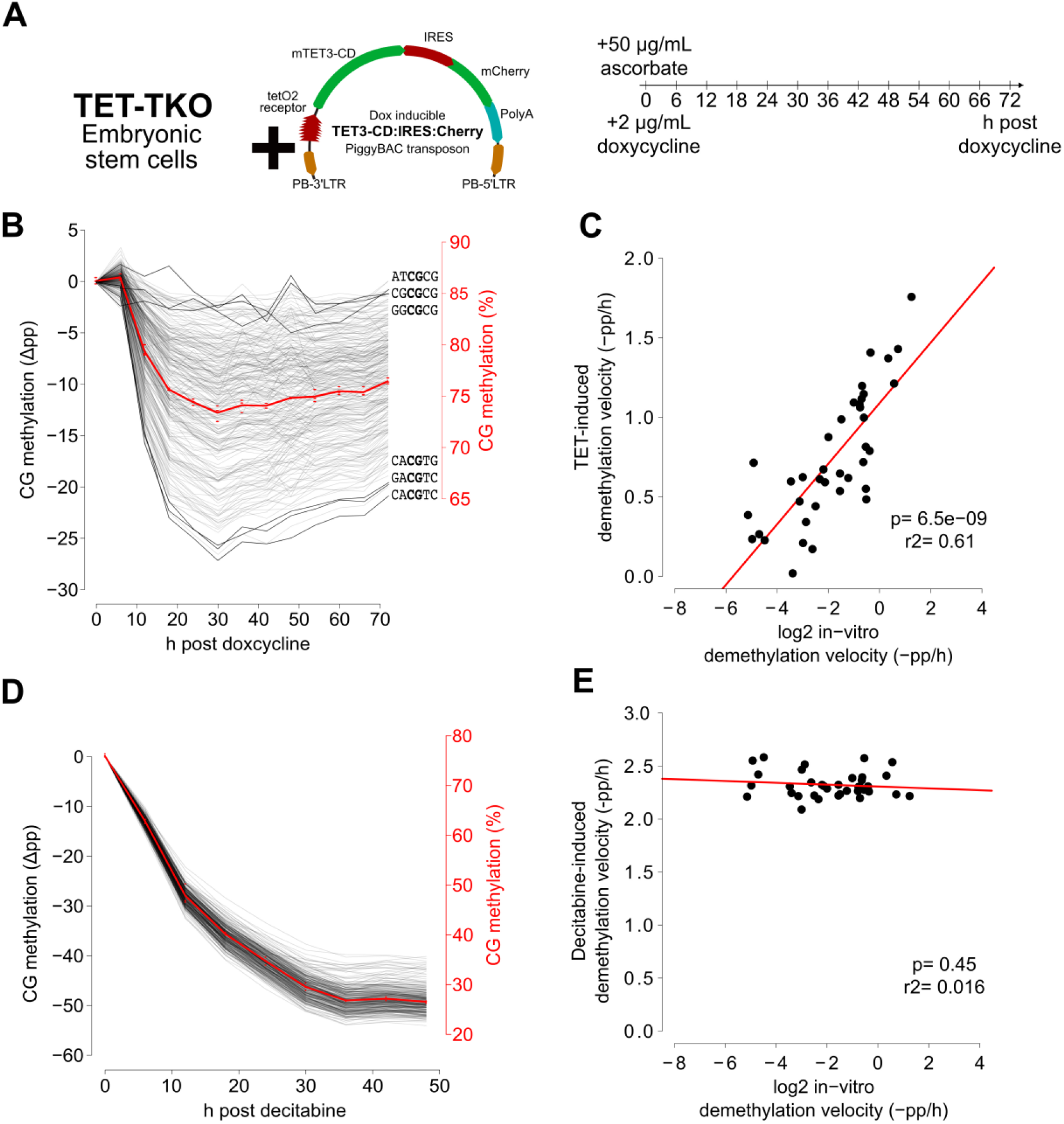
TET3 catalytic domain selectivity in cultured mouse embryonic stem cells is proportional to selectivity observed *in vitro*. **(A)** The TET3 catalytic domain (TET3-CD) was overexpressed in a TET triple-knockout (TET-TKO) embryonic stem cell line using an inducible piggyBAC transposon system. Doxycycline treated (Dox) and control (no-Dox) samples were collected, in triplicate, every 6 h over a 72 h period. **(B)** Absolute methylation levels in non-CGI contexts (red) and the difference in methylation of individual motifs (grey lines) following Dox-induced TET3-CD expression. **(C)** Demethylation velocities of CG-containing hexamer motifs from B (calculated from linear phase, 6-18 h), compared to log2 demethylation velocity *in vitro* (Figure 1). **(D)** Absolute methylation levels in non-CGI contexts (red) and the difference in methylation of individual motifs (grey lines) following demethylation by the small molecule Decitabine. **(E)** Demethylation velocities of CG-containing hexamer motifs from E (calculated from linear phase, 0-12 h), compared to log2 demethylation velocity *in vitro* (Figure 1).

To determine whether TET3 displayed any sequence specificity within this system, we binned each mapped CG dinucleotide according to the 2 base pairs flanking it in each direction. Inspection of individual CG-containing hexamer sites revealed a large variation in the demethylation velocity between motifs. We predicted that at least some of this variation in rate was due to low starting methylation of some motifs (49 motifs had <65% of starting methylation, Figure S6C, left panel). These lowly-methylated motifs often contained CGs that were in addition to the central CG (red dots), indicating that they were likely restricted to CpG island regions (CGI) which are well-known for significantly reduced methylation levels. Indeed, when we considered only motifs located in non-CGI regions, starting methylation for all 256 motifs was much more consistent at 75.8-90.2 % (Figure S6C, right panel) and was thus used for further analysis.

Most CG-containing hexamer motifs showed linear demethylation 6-18 h post dox treatment, allowing calculation of maximum demethylation velocity for each motif and comparison to the 38 CG-containing motifs from the *in vitro* experiment (Figure 4C). Despite a much larger range of demethylation velocities observed in the *in vitro* experiment, a significant correlation was observed between the demethylation velocities in the cell culture experiment and log2 transformed values from the synthetic *in vitro* data (r2 = 0.61, p = 6.5e-9). This indicates that the unique selectivity of TET observed in *in vitro* biochemical reaction also exists in a cellular context.

To be more confident that the observed variation in mTET3-targeting was not a technical artifact, we examined motif demethylation dynamics in wild-type V6.5 mESCs following treatment with the demethylating small molecule decitabine (Figure 4D). Due to the fact that decitabine drives demethylation by inhibition of DNMT1 (*19*) (and thus operates in a TET-independent manner), we hypothesized that similar methylation site preference should not exist. Indeed, overall there was minimal variation in motif-demethylation rate following decitabine treatment (Figure 4D), but most importantly, when compared to the *in vitro* demethylation, no significant correlation in demethylation velocity was uncovered (Figure 4E) (r2 = 0.016, p = 0.45).

To further characterize mTET3 selectivity in mESCs cells, we investigated which CG sites were demethylated the fastest and slowest according to the nucleotides at each flanking position when all other bases were kept the same (Figure 5A, top panel); a measure we termed intra-motif positional preference. In doing so, we discovered that for 95.3% of motifs (i.e. 61/64), those with adenine immediately upstream of CG (i.e. −1 position) were demethylated faster than any other motif with base in that position. In the 3 remaining cases, the preferred base at −1 was C – a result perfectly matching our expectations from our *in vitro* data as well as the structural analysis and modelling. Moreover, when the +1 position was examined, for 79.6% of motifs (51/64), those with T demethylated faster than another base, with A being the next most common (Figure 5A, bottom panel). Favoured nucleotides at the –2 and +2 position were less obvious; C and G containing motifs were the most rapid demethylating in 54.6% (35/64) and 40.6% (25/64) of instances, respectively. Remarkably, the most preferred nucleotides at each position recapitulated the CACGTG E-box motif (Figure 5B), as initially uncovered in the *in vitro* experiments.

**Figure 5.**
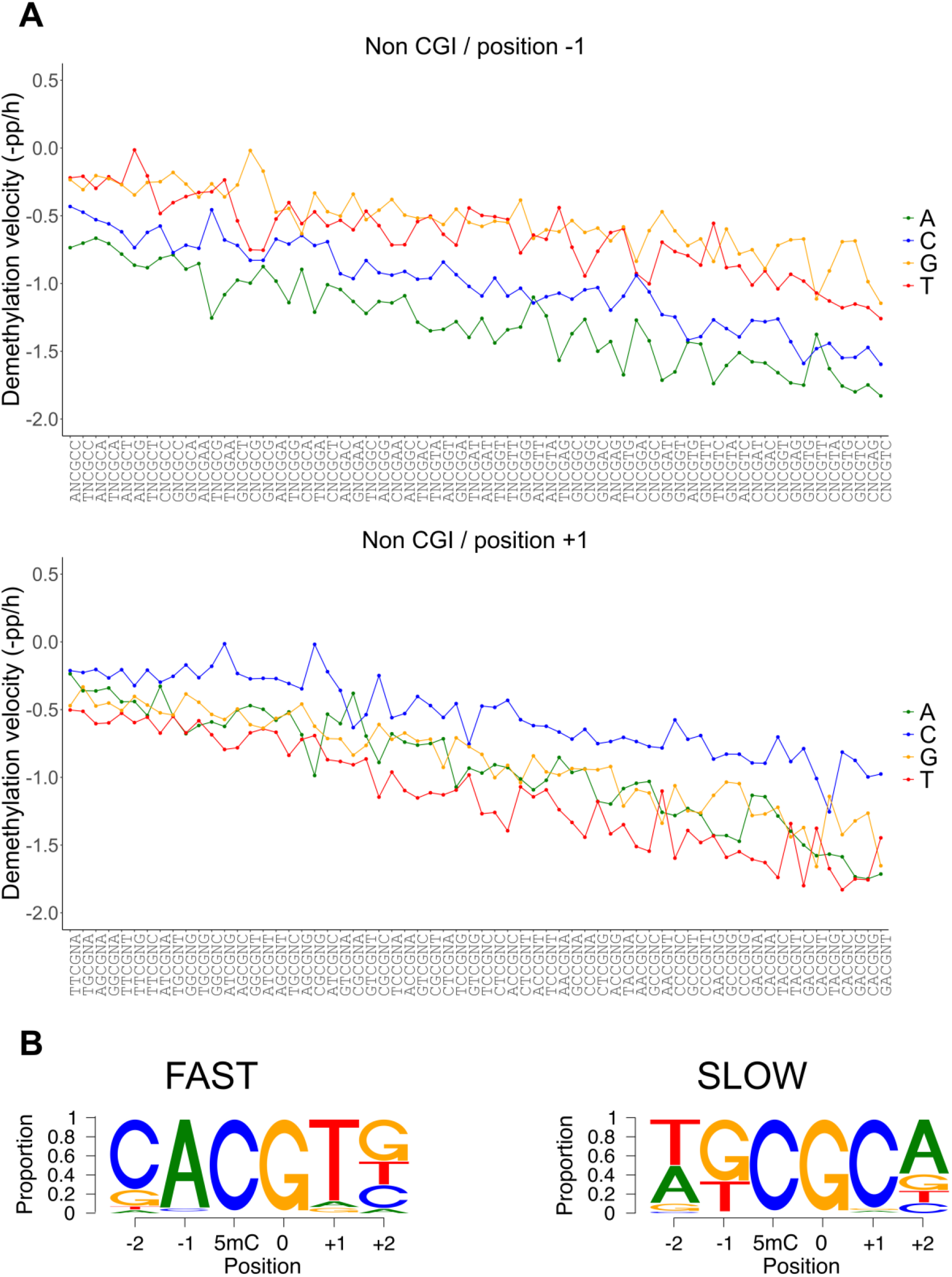
Intra-motif positional preference (IMPP) for TET3-CD induced demethylation in cultured embryonic stem cells. **(A)** Demethylation kinetics of the 4 nucleotides in a given CG containing hexamer position (shown are −1 and +1 positions flanking the central CG). **(B)** Proportion of nucleotides which are demethylated the fastest (left) and slowest (right) at each motif position when all other nucleotides in that motif are kept constant.

When demethylation rates of each of the 256 CG-containing hexamers were considered individually, the CACGTG E-box sequence was the third most preferentially targeted motif (−1.76% per hour) (Figure 6A). As mentioned, E-box is notable for binding c-MYC, an iconic ‘immediate-early’ bHLH- and bZIP-domain-containing protein that is amongst the first to be transcribed in response to a wide variety of cellular stimuli. Significantly, the two motifs that demethylated faster (CACGTC and GACGTC) also constitute binding sites for bZIP-domain, methylation-sensitive ‘immediate-early’ transcription factors, i.e., CREB and JUN/FOS, respectively. A further 5 motifs in the top 15 favorable recognition sites bind to bZIP or bHLH domain-containing transcription factors, many of which display methylation-sensitive binding (*20*) (Figure 6A). In contrast, the least preferred bases at each position most commonly featured G and C at the –1 and +1 position, with the TGCGCA OCT4 binding motif the least preferred (Figure 6A, lower panel). Interestingly, OCT4 and other members of the POU transcription factor family are known to bind TGCGCA specifically when methylated (*20*), indicating that resistance to TET demethylation may be a prerequisite for OCT4 binding.

**Figure 6.**
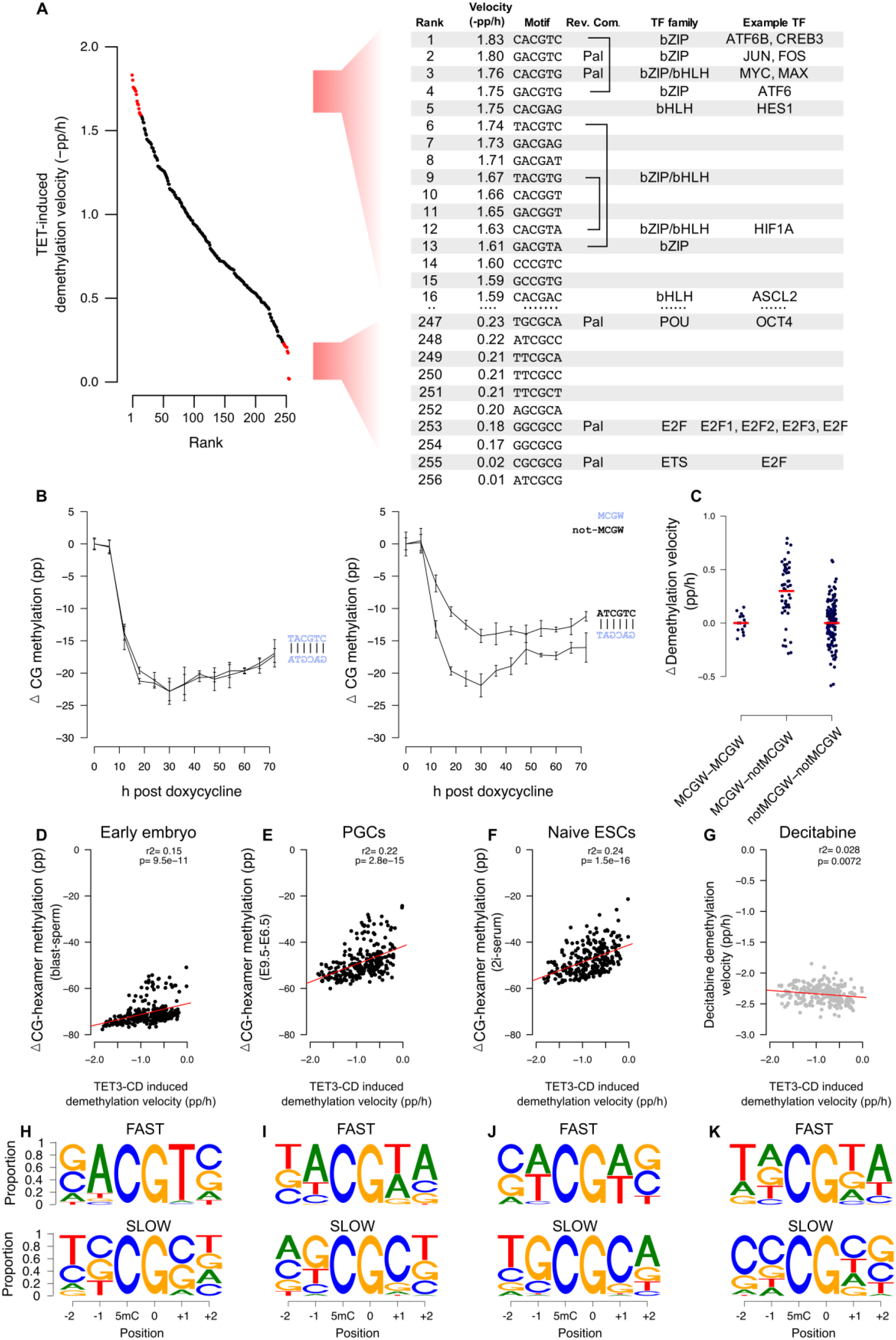
TET selectivity in cultured cells coincides with methylation-sensitive transcription factors, is strand dependent, and correlates with global demethylation *in vivo*. **(A)** Many fast demethylating CG-containing hexamer motifs following TET3-CD expression (rank 1-16) bind bZIP and bHLH methylation-sensitive transcription factors. Slow demethylating sites (rank 247-256) bind methylation-sensitive E2F and POU family transcription factors. **(B)** Many complementary motifs have similar demethylation kinetics (e.g. TACGTC, left panel); however, some are significantly discordant (e.g. ATCGTC, right panel). **(C)** CG-containing hexamers with MCGW on one strand demethylate faster than those without MCGW on the complementary strand (MCGW-notNCGW). In contrast, motifs where MCGW is present on both strands (MCGW-MCGW) or not (notMCGW-notMCGW) show equal demethylation rates. **(D-G)** CG-containing hexamer demethylation velocities following TET3-CD overexpression (x-axis, TET3-CD induced demethylation velocity) and the demethylation found in (y-axis) mouse **(D)** embryos following fertilization (blast-sperm), **(E)** primordial germ cells (E9.5-E6.5), **(F)** naive embryonic stem cells. Only a weak negative correlation exists with **(G)** decitabine-treated cells. (**H-K**) Intra-motif positional preference (IMPP) for the demethylating scenarios listed in D-G. Shown is the proportion of nucleotides that are demethylated the fastest (upper panel) and slowest (lower panel) at each motif position when all other nucleotides in that motif are kept constant.

Many rapidly demethylating non-palindromic sites showed demethylation velocities on each strand that were similar. For example, of the 10 fastest demethylating CG-containing hexamers, 3 had reverse-complement motifs also in the top-20 (Figure 6A), implying that both strands of a CG-containing hexamer may be targeted with similar efficiency. Despite this, we uncovered striking exceptions, where certain CG-containing hexamer sequences were much preferred over their antisense counterparts (Figure 6B). We noticed that preferred motifs in a discordant antisense pair were often MCGW, whereas the non-favored strand was non-MCGW (Figure 6C). In addition to confirming our biochemical and structural predictions, this result is significant because it shows demethylation rates on each strand are clearly separable in situations of discordant binding preference. As such, we conclude they are likely demethylated by independent binding events.

Mammalian DNA is subject to two significant waves of demethylation during normal development. The first erasure event occurs in the zygote immediately following fertilization (*21*), while the second occurs during primordial germ cell specification and proliferation (*21*). Given that zygotic methylation erasure occurs without cell replication, a large component of the demethylation must be active and driven by the oocyte-specific ‘TET3o’ variant that lacks the CXXC domain and resembles the construct we used (*7*). The kinetics of demethylation in mouse primordial germ cells implies that demethylation is largely a passive process; however, TET enzymes play an active role in methylation erasure during this time as well (*22*).

To test whether the intrinsic sequence preferences of the TET catalytic domain could be detected during global methylation erasure in post-fertilization embryos, primordial germ cells and naïve stem cells, we re-examined the published BS-seq data. While these studies did not sample throughout the demethylation time-course as we did in our mTET3-CD overexpression experiments (Figure 4A), we still found significant correlations between all global demethylation experiments tested and our results (Figure 6D-F), particularly when lowly methylated CGI-rich motifs were removed. Moreover, when we analyzed the intra-motif positional preference in these datasets, we found that all 3 datasets recapitulated preference for A at –1, and A or T at +1 positions (Figure 6H-K, upper panels). Additionally, we found that when de novo methyltransferases were removed from 3 independent human embryonic stem cells (*14*), the extent of demethylation following unopposed TET activity was correlated with the CG-containing hexamer demethylation rate in our overexpression model (r2>0.43, p<7e-33, n=3; Figure S7A). Likewise, the CG-containing hexamer demethylation rates from our cell culture experiment were significantly correlated with the average predicted ‘TET activity’ on 800,000 CpG sites in mouse ESCs from in a previous study (*23*) (r2=0.29, p=8.9e-21, Figure S7B).

To exclude the possibility that some other factor was driving this relationship in CG-containing hexamer demethylation rate (e.g., chromatin structure), we assessed TET-independent global demethylation driven by decitabine treatment and found no correlation (Figure 6G). Moreover, intra-motif positional preference for those sites losing methylation the fastest following decitabine treatment did not feature any selectivity at –1 or +1 (where we find TET favored A and T in the strict sense, or more loosely M and W), but instead we uncovered a previously described DNMT1 motif preference, TNCGNW (*24*) (Figure 6K).

Together, our data demonstrate that the TET catalytic domains possess a previously unknown intrinsic sequence specificity that orchestrates DNA demethylation and can be detected in a wide range of methylation erasure scenarios, both *in vivo* and in culture. We show that in addition to other identified targeting mechanisms, in particular CXXC domain targeting and chromatin factors, the intrinsic sequence preference of the TET catalytic domains significantly contributes to the establishment of DNA methylation patterns and TET function in the cell. Highly favored motifs constitute immediate-early transcription factors, which are the first to respond to a range of stimuli such as mitogens or infection and proceed to initiate expression of downstream effector genes by binding to DNA in a methylation-sensitive manner. Thus, it makes biological sense that their binding sites should be efficiently targeted for DNA methylation erasure. Furthermore, where TET apparently acts to remove DNA methylation from DNA to allow binding of immediate-early effectors, it also may help preserve DNA methylation at OCT4 sites (by avoiding them) to equally stimulate binding and maintenance of developmental potency (Figure S8). Together, our data support a model where on multiple levels, the kinetics of TET-mediated demethylation is inextricably tied to the biological function of the TET enzymes.

## Supporting information

Combined Supplementary Information

## Acknowledgments

We thank the High Throughput Sequencing unit of the Genomics & Proteomics Core Facility, German Cancer Research Center (DKFZ), for providing excellent next-generation sequencing services. We are grateful to Dr hab. Krzysztof Skowronek (IIMCB) and Dr Radosław Pluta (IIMCB) for the technical advice and assistance in the project. We want to thank Prof. Yanhui Xu (Fudan University) for sharing the hTET2 expression construct used for crystallization. Diffraction data for the hTET2:DNA complexes have been collected on BL14.1 at the BESSY II electron storage ring (Helmholtz-Zentrum Berlin) and the DESY P11 beamline (PETRA III, Hamburg). We want to acknowledge Dr Manfred Weiss (Helmholtz-Zentrum Berlin) and Dr Anja Burkhardt (DESY) for their kind assistance during the diffraction data collection.

## Funding

University of Otago (TAH)

Deutsche Forschungsgemeinschaft JU2773/2-1 (TPJ)

Foundation for Polish Science POIR.04.04.00-00-5D81/17-00 (MB)

Polish National Science Centre 2018/30/Q/NZ2/00669 (MB)

Polish National Agency for Academic Exchange PPI/APM/2018/1/00034 (MB)

Deutsche Forschungsgemeinschaft ZA153/28-1 (MZ)

## Author contributions

Conceptualization: TPJ, TAH, MB

Methodology: MR, DR, OOR, CID, CRG, AHW, MW, MRazew, IMMayyas, KM, US, OK, KZ, IMMorison, DW, FK, TPJ, TAH, MB

Investigation: MR, DR, OOR, CID, CRG, AK, KM, OK, US, KZ, TPJ, TAH, MB

Visualization: MR, DR, OOR, CID, CRG, US, RZJ, FK, TPJ, TAH, MB

Funding acquisition: MZ, TPJ, TAH, MB

Project administration: TPJ, TAH, MB

Supervision: TPJ, TAH, MB, CP, MZ, RZJ, MH

Writing – original draft: TPJ, TAH, MB

Writing – review & editing: All authors

## Data and materials availability

All biological materials are available on request. The assembled construct sequence for pB-tetO2-mTET3cd-mCherry can be found on the NCBI sequence archive under accession (MW139646). In-house developed analysis scripts and processed data are available on GitHub. (https://github.com/TimHore-Otago/TET_specificity). Raw high-throughput sequencing data is available on GEO under accession GSE159205. *Note to reviewers: use the following token to access this data:* **irkpckachtmhvuv**

## Supplementary Materials

Materials and Methods

Table S1 – S2

Fig S1 – S8

References (25 – 40)

## References and Notes

1. M. G. Goll, T. H. Bestor, Annu. Rev. Biochem. 74, 481–514 (2005).

2. M. Tahiliani et al., Science. 324, 930–935 (2009).

3. Y.-F. He et al., Science. 333, 1303–1307 (2011).

4. Y. Costa et al., Nature. 495, 370–374 (2013).

5. C. A. Doege et al., Nature. 488, 652–655 (2012).

6. T. A. Hore et al., Proc. Natl. Acad. Sci. U.S.A. 113, 12202–12207 (2016).

7. T.-P. Gu et al., Nature. 477, 606–610 (2011).

8. O. Abdel-Wahab et al., Blood. 114, 144–147 (2009).

9. M. Bochtler, A. Kolano, G.-L. Xu, Bioessays. 39, 1–13 (2017).

10. H. Hashimoto et al., Nucleic Acids Res. 40, 4841–4849 (2012).

11. M. Ko et al., Nature. 497, 122–126 (2013).

12. T. Nakagawa et al., Molecular Cell. 57, 247–260 (2015).

13. K. Williams et al., Nature. 473, 343–8 (2011).

14. J. Charlton et al., Nat. Genet. 52, 819–827 (2020).

15. H. Hashimoto et al., Nature. 506, 391–395 (2014).

16. L. Hu et al., Cell. 155, 1545–1555 (2013).

17. L. Hu et al., Nature. 527, 118–122 (2015).

18. X. Hu et al., Cell Stem Cell. 14, 512–522 (2014).

19. P. A. Jones, S. M. Taylor, Cell. 20, 85–93 (1980).

20. Y. Yin et al., Science. 356 (2017), doi:10.1126/science.aaj2239.

21. W. Reik, W. Dean, J. Walter, Science. 293, 1089–1093 (2001).

22. J. A. Hackett et al., Science. 339, 448–452 (2013).

23. P. A. Ginno et al., Nat Commun. 11, 2680 (2020).

24. S. Adam et al., Nat Commun. 11 (2020), doi:10.1038/s41467-020-17531-8.

