## Supplementary material for "Pronounced sequence specificity of the TET enzyme catalytic domain guides its cellular function": Combined Supplementary Information

#### Materials and Methods

##### Cloning, expression, and purification of TET enzymes

Construction of the pET28a vectors containing the catalytic domains of mouse TET1, TET2, TET3 has been described previously (*1*). Full-length *Naegleria gruberi* TET (nTET) was cloned from a synthetic gene (IDT) into pET28a bacterial expression vector in fusion with the N-terminal His-tag. The pET28a vectors encoding each TET catalytic domain were transformed in *E. coli* BL21 (DE3) CodonPlus RIL (Novagen) and grown on kanamycin and chloramphenicol selection plates overnight. For protein expression, single colonies were inoculated in 100 ml of LB media supplemented with appropriate antibiotics and grown overnight at 37 °C in a shaking incubator. For each protein expression, 4 L of LB media were inoculated with the overnight culture and the cells were grown at 37 °C until OD600 0.6 was reached, recombinant protein expression was induced with 0.5 mM IPTG and culture was grown at 20 °C for ~14-15 hours. The cells were harvested by centrifugation (Lynx 6000 (Thermo), Fiberlite F9-6 1000 LEX, at 4200 RPM for 15 min), washed with 1x STE buffer (10 mM Tris-HCl pH 8.0, 100 mM NaCl, 1 mM EDTA) and stored at –20 °C until further use. For purification, cells were resuspended in lysis buffer (50 mM HEPES pH 6.8, 35 mM imidazole, 1 mM α-ketoglutarate, 1 mM DTT, 500 mM NaCl, 10% glycerol, supplemented with protease inhibitor cocktail (Roche)) and disrupted using Bandelin Sonoplus ultrasonic homogenizer. The cell lysates were cleared by centrifugation (Lynx 600 (Thermo), Fiberlite F21-8x50y) at 38,300 xg for 75 minutes at 4 °C and the supernatant was loaded onto an affinity column containing 2 ml of Ni-NTA agarose beads (Genaxxon, Germany). The beads were washed thoroughly with wash buffer (50 mM HEPES pH 6.8, 35 mM imidazole, 1 mM α-ketoglutarate, 1 mM DTT, 500 mM NaCl, 10% glycerol) and eluted with elution buffer (50 mM HEPES pH 6.8, 300 mM imidazole, 1 mM DTT, 500 mM NaCl, 10% glycerol). The concentrated fractions were pooled and dialyzed against dialysis buffer (50 mM HEPES pH 6.8, 1 mM α-ketoglutarate, 1 mM DTT, 300 mM NaCl, 10% glycerol) and subsequently aliquoted, flash-frozen in liquid nitrogen and stored at –80 °C until further use.

#### Preparation of TET *in vitro* substrates

The substrates for *in vitro* reactions: human CG-rich (249 bp), promotor fragments of mouse Esrrb (249 bp), mouse Nanog (260 bp), and mouse Tcl1 (251 bp) were amplified from genomic DNA, subcloned into pCR2.1 vector using the TOPO TA Cloning kit (Invitrogen) and sequence confirmed (substrate sequences are listed in Table S2). Fully 5mC-modified substrates with methylated cytosine in both CG and non-CG contexts were amplified from subcloned template plasmids using primers listed in Table S2. Fully 5mC-modified substrates with methylated cytosine in both CG and non-CG contexts were amplified by PCR from subcloned template plasmids (primers listed in Table S2) using in-house-made Taq polymerase and dNTP mixture containing 5mdCTP (NEB, N0356S) instead of dCTP. To generate substrates only methylated at CpG sites, unmethylated templates were amplified by PCR using in house made TaqPol with standard dNTP mix and the purified PCR products were further methylated with M.SssI methyltransferase (NEB, M0226S) following the manufacturer's protocol. The amplified substrates were purified using #4.1 BOMB protocol (*2*) and verified with gel electrophoresis.

For LC-MS, 26 bp hemimethylated substrate was prepared by annealing complementary oligonucleotides (IDT, listed in Table S2). Briefly, 20 µM of each oligo was resuspended in annealing buffer (10 mM HEPES pH 7.4 and 50 mM NaCl). The oligos were heated at 85°C for 5 minutes and were cooled down gradually to room temperature over several hours. The annealed oligos were stored at –20 °C until further use.

#### TET activity assays

To analyze the activity of mammalian TETs, 0.15 µM DNA substrate was incubated with 2 µM enzymes in a reaction mixture containing 50 mM HEPES pH 6.8, 100 µM $Fe^{2+}$, 1 mM, α-ketoglutarate, 1 mM ascorbic acid, 150 mM NaCl at 37 °C for up to 120 minutes. The reaction was stopped at specified time points by adding an aliquot of the reaction mixture to 1% SDS and proteinase K (NEB) for an hour at 50 °C. Proteinase K was inactivated and the DNA was purified using DNA purification kit (Macherey-Nagel) following the manufacturer's protocol. The purified DNA was ligated with methylated TruSeq LT Illumina adapters and subjected to bisulfite conversion using EZ-DNA Methylation-Lightning kit (Zymo Research, D5030) according to the manufacturer's protocol. The bisulfite converted DNA was amplified by PCR (primer details in Table S2) and successful amplification was verified using gel electrophoresis. The amplified products were quantified using NEBNext kit (E7630S, NEB) and sequenced on an Illumina MiSeq 2x300 bp platform (Illumina).

For the nTET activity assay, the reaction was carried out as outlined above with slight modifications. Briefly, 0.5 µM DNA substrate was treated with 2 µM enzyme in a reaction mixture containing 50 mM Bis-Tris (Sigma-Aldrich, 14879-100G-F), pH 6.0, 75 µM $Fe^{2+}$, 1 mM α-ketoglutarate (Sigma-Aldrich, 75890-25G), 1 mM ascorbic acid and 100 mM NaCl for specified time at 34 °C. After enzymatic treatment, the DNA was processed using same procedure as described above for the mammalian enzymes.

#### Liquid chromatography mass spectrometry (LC-MS)

To quantify the oxidized products of TET enzyme using LC-MS, 0.5 µM of hemimethylated 26 bp dsDNA substrates (oligonucleotide sequences listed in Table S2) were treated with mTET3 CD for 1, 5, 10, 20 and 40 minutes. The reactions were stopped at specified time points by adding 10 mM EDTA and flash frozen in liquid nitrogen. Subsequently, the enzyme was inactivated at 95 °C for 5 minutes, followed by proteinase K (New England Biolabs, P8107S) treatment for an hour at 50 °C. The DNA was recovered by ethanol precipitation and analyzed by LC-MS.

For this, the purified DNA oligonucleotides were digested into nucleosides using the following protocol: 160 ng of each DNA, 0.6 U nuclease P1 from *Penicillium citrinum* (Sigma-Aldrich), 0.2 U snake venom phosphodiesterase from Crotalus adamanteus (Worthington), 200 ng pentostatin (Sigma-Aldrich) and 500 ng tetrahydrouridine (Merck-Millipore) were incubated at 37 °C in 5 mM ammonium acetate pH 5.3 (Sigma-Aldrich) for 2 h. Afterward, 1/10 vol. of 10x fast alkaline phosphatase buffer (50 mM NH4OAc pH 9.0) and 1 U fast alkaline phosphatase (Fermentas) were added, followed by incubation at 37 °C for 1 h. The nucleosides were then spiked with labeled internal standard (D3-dm5C and D2-hdm5C) and subjected to analysis. For each time point, 400 fmol digested DNA oligo and 100 fmol of each internal standard were injected and analyzed via LC-MS (Agilent 1260 series and Agilent 6460 Triple Quadrupole mass spectrometer equipped with an electrospray ion source (ESI)). The solvents consisted of 5 mM ammonium acetate buffer (pH 5.3; solvent A) and LC-MS grade acetonitrile (solvent B; Honeywell). The elution started with 100% solvent A with a flow rate of 0.35 ml/min, followed by a linear gradient to 20% solvent B at 10 minutes and back to 100% solvent A in 2 minutes. Initial conditions were regenerated with 100% solvent A for 5 minutes. The column used was a Synergi Fusion (4 µM particle size, 80 Å pore size, 250 × 2.0 mm; Phenomenex). The UV signal at 254 nm was recorded via a diode array detector (DAD) to monitor the main nucleosides. ESI parameters were as follows: gas temperature 350°C, gas flow 8 l/min, nebulizer pressure 50 psi, sheath gas temperature 350°C, sheath gas flow 12 l/min, capillary voltage 3000 V. The MS was operated in the positive ion mode using Agilent MassHunter software in the dynamic MRM (multiple reaction monitoring) mode. For quantification, a combination of external and internal calibration was applied as described previously (*3*).

##### Bioinformatic analysis of sequencing from *in vitro* demethylation

The quality control of the next generation sequences (NGS) in FASTQ format was done using FastQC (Babraham Bioinformatics). The adapters from the sequences were trimmed and quality controlled using TrimGalore (Babraham Bioinformatics). Files were then demultiplexed using a custom script and processed further using BiQ Analyzer HT (Max-Plank-Institut Informatik), where all Cs in CpN contexts were analyzed and exported to an Excel readable file. The average oxidation at each C was calculated and the overall oxidation for each time point is obtained by subtracting the oxidation at respective time point from the control. Linear regression was used to estimate the demethylation velocity for each site.

##### Linear regression model to predict reaction rates

For quantitative analysis of the sequence dependence of demethylation rates, we assumed uncorrelated sequence preferences. Under this assumption, the logarithms of reaction rates should be amenable to linear regression (in the R language) with DNA base identities as categorical variables. Two bases upstream of the C, and two bases downstream of the C were included in all models. For the *in vitro* data, the position of the G was also kept variable. For the *in vivo* data, a G in this position was assumed to have enough statistics for well-defined slopes. For the *in vitro* data, two separate slope determinations were available. The absolute differences (augmented by 0.1 to avoid overemphasis on data points with accidentally good agreement) were used as weights for the regression analysis. To exclude overfitting, data were split into “training” and “testing” sets. Except for the TET1 *in vitro* data, correlation coefficients were very similar when the full data set was used for training and testing and when training and testing data were non-overlapping.

##### Expression of human TET2 for structural studies

For structural studies, we used a fragment of human TET2 (1129–1936) with residues 1481–1843 replaced by a 15-residue GS linker. The protein was expressed with an N-terminal $His_6$-$Gly_3$-$His_6$-SUMO-tag. This was done in E. coli BL21 (DE3) CodonPlus RIL cells (Novagen) under T7 promoter control from a pET28a derived plasmid, which was maintained under kanamycin and chloramphenicol selection. A pre-culture at OD=0.8 1/cm was induced with 0.1 mM isopropyl β-d-1-thiogalactopyranoside (IPTG) and grown overnight at 16 °C (shaking at 130 RPM). Cells were harvested by centrifugation (4000 g at 4 °C for 30 min) and bacterial pellets were frozen and stored at -20 °C until use.

##### Protein purification of human TET2 for structural studies

The hTET2 bacterial pellets from 2 L of culture were resuspended in 50 ml Sonication buffer (50 mM HEPES, pH 8.0, 500 mM NaCl, 10 mM imidazole, 20% glycerol, 5 mM β-mercaptoethanol, 2 mM PMSF) and sonicated on ice (15 s of pulse, 45 s of rest, 5.5 power, total sonication time 6 min). The lysate was cleared by ultracentrifugation (4 °C, 40 min, 40 000 g), and applied two times on a column containing 5 ml of Ni-NTA resin (Qiagen) equilibrated with the Sonication buffer. The column was washed sequentially with 100 ml Wash1 buffer (20 mM HEPES, pH 8.0, 500 mM NaCl, 10 mM Imidazole, 5 mM β-mercaptoethanol), 500 ml Wash buffer (20mM HEPES, pH 8.0, 1750 mM NaCl, 1 mM imidazole, 5 mM β-mercaptoethanol), DnaK buffer to remove DnaK chaperone (20 mM HEPES, pH 8.0, 50 mM NaCl, 2 mM MgCl2, 2 mM ATP, 5 mM β-mercaptoethanol) and Ulp1 buffer (20 mM HEPES pH 8.0, 150 mM NaCl, 15 mM imidazole, 5 mM β-mercaptoethanol). The protein was cleaved from the tag on-column (170 µg of Ulp1 in Ulp1 buffer for protein from 10 L of bacterial culture, 15h, 6 °C). Eluted protein was diluted in Dilution buffer (20 mM HEPES, pH 8.0, 5 mM β-mercaptoethanol), and applied to a 5ml Heparin column (GE Healthcare) equilibrated in Hep1 buffer (20 mM HEPES, pH 8.0, 40 mM NaCl, 5mM β-mercaptoethanol), and eluted with a gradient between this buffer and Hep2 buffer (20 mM HEPES, pH 8.0, 2000 mM NaCl, 5mM β-mercaptoethanol). hTET2 fractions were then concentrated to 1 ml and subjected to gel filtration on a Superdex 200 (GE Healthcare) column equilibrated in GF buffer (10 mM HEPES, pH= 7.4, 100 mM NaCl and 1 mM DTT).

##### Crystallization and structure determination

For crystallization, we used equimolar mixtures of hTET2 (from a 70 mg/ml stock in GF buffer) and 12-mer dsDNA (from IDT, in water). Top strand sequences of most and least optimal substrates were 5’-ACACA5mCGTGTGT 3’ and 5’-ACAGG5mCGCCTGT-3’, respectively. Complementary bottom strands, also with DNA methylation, were annealed into top strands by heating to 95 °C for 10 min and subsequent slow-cooling. For crystallization, 1.5 µl of an 0.5 mM solution of hTET2-dsDNA complex were mixed with 1.5 µl of Reservoir buffer (100 mM MES, pH 6.3, PEG 2000 monomethyl ether), and supplemented with 2 mM of the co-substrate analog N-oxalylglycine (NOG) and 1 mM $Fe^{2+}$ (favorable substrate) or $Mn^{2+}$ (unfavorable substrate).

Crystals were grown in hanging drops at 4 °C by equilibration of the crystallization mix against Reservoir buffer. For the complex of hTET2 with the least optimal substrate, seeding was required. A crystal of insufficient quality for diffraction experiments was crushed, crystal seeds were suspended in 50 µl reservoir buffer, and this stock was then used for seeding (seed solution was 10% of the crystallization drop volume. Diffraction data up to 2.0 Å resolution for the complexes with most and least optimal substrates were collected at beamlines of the DESY (P11, DESY, Hamburg) and BESSY (Berlin), respectively, at 100 K. Both datasets were processed with XDSapp (*4*). Structures were solved by the Phaser program (*5*), using the previous PDB TET2 model (PDB accession: 4NM6) as template, and refined using Phenix software (*6*) (Table S1).

#### Molecular dynamics simulations

All molecular dynamics (MD) and energy minimization were performed using the Amber16 simulation package (*7*). Using the PDB:4NM6 structure as reference the hexameric recognition sequence AC5mCGGT (in PDB accession: 4NM6) was systematically varied at positions 2 and 5 (the positions flanking the CpG) to generate all 16 possible sequence variants using the xleap module of Amber16. The resulting complex structures were conformationally relaxed using energy minimization (5000 conjugated gradient steps), short MD simulations at 290 K for 1 ns followed by another energy minimization (5000 steps). For the protein, the parm14SB and for the DNA the parmBsc1 force field were used in combination with a Generalized Born implicit solvent as implemented in Amber16 (igb=5 option). During each minimization and MD phase the backbone atoms of the DNA and of the protein were restrained to the positions found in the reference structure (PDB accession: 4NM6) but allowing full mobility of the side chains.

#### Embryonic Stem Cell Lines and Culturing Conditions

Mouse embryonic stem cell culture was performed using standard media conditions in the presence of serum and leukemia inhibitory factor (LIF) (DMEM, 4,500 mg/l glucose, 4 mM L-Glutamine and 110 mg/l sodium pyruvate, 15% fetal bovine serum, 1 U/ml penicillin 1 mg/ml streptomycin, 0.1 mM non-essential amino acids, 50 mM β-mercaptoethanol, 1000 U/ml LIF). All cells were grown 'feeder-free' on gelatin-coated plates and were maintained at 37 °C in a humidified 5% $CO_2$-containing atmosphere.

To create an embryonic stem cell line with inducible TET expression, we first created a PiggyBAC construct that contained the mouse TET3 catalytic domain (mTET3-CD) under the control of a tetracycline responsive promoter (pB-tetO2-mTET3cd-mCherry). The correct identity of this construct was confirmed using shotgun Illumina sequencing on the MiSeq platform. The assembled construct sequence can be found on the NCBI sequence archive under accession (MW139646).

A construct mix containing 4 µg of pB-tetO2-mTET3cd-mCherry, 4 µg of pB-CAG-rtTA-Puro (containing the reverse tetracycline-controlled transactivator (rtTA) gene) and 4 µg of the pCAG-pBASE vector (containing a PiggyBac transposase allowing "cut and paste" insertion of the construct into the host genome (*8*)) was prepared in 125 ml of Opti-MEM® Reduced Serum Media (Thermo Fisher Scientific, cat. 31985062). Transfection Reagent was prepared separately by adding 89 µl of Opti-MEM® to 36 µl of FuGENE® HD (Promega, E2311) then incubated for 5 minutes. Once the construct mix and FuGENE® solutions were incubated separately for 5 minutes, they were added together and incubated for an additional 20 minutes before being added dropwise to TET triple-knockout ESCs, thus following a previously established protocol (*9*). Cell cultures were then incubated at 37°C for 2 to 6 hours in 250 µl of media allowing transfection to take place. Selection for positive transfectants was performed over 7 days by adding puromycin to the media (2 µg/ml).

Once a stable mTET3-CD complemented TET-TKO line was produced, media supplemented with 50 µg/ml Ascorbate (Sigma, A7631) was added to cells plated at low-density in 12-well plates overnight. The following morning, mTET3-CD expression was initiated by the addition of 1 µM doxycycline. Control and doxycycline-treated cells were harvested in triplicate every 6 h over the next 72 h.

For the decitabine treatment experiment, wild-type V6.5 hybrid embryonic stem cells (i.e., C57BL/6 X 129/sv cross; a gift from Bjorn Oback) were seeded at low-density, treated with 0.215 µM of decitabine (5-aza-2'-deoxycytidine) over a period of 48 hours. This level of decitabine had been determined empirically in a prior experiment as being the highest concentration that did not cause overt cell death. In both experiments, cells were harvested by media removal and addition of a 4 M guanidinium isothiocyanate-based lysis buffer (GITC) (*2*). Cell lysates were then stored at -80 °C ahead of total nucleic acid purification using the Bio-On-Magnetic-Beads (BOMB) system (*2*). Briefly, cell lysate was combined with TE-diluted Sera-Mag Magnetic SpeedBeads (GE Healthcare, GEHE45152105050250) and isopropanol in a volumetric ratio of 2:3:4 (beads:lysate:isopropanol). Beads were captured with a neodymium magnet and washed once with isopropanol, twice with 70% ethanol and resuspended in milliQ water.

**PBAT library preparation and sequencing**

Bisulfite-converted genomic libraries were prepared using a modified post-bisulfite adaptor tagging (PBAT) method (*10*). Purified DNA was subjected to bisulfite conversion using the EZ-96 DNA methylation Direct™ MagPrep kit (Zymo Research, D5044), according to the manufacturer's user guide, however, reagent volumes were scaled down to 25% of the recommended volume, with the exception of the final elution step which remained at 25 µl. To synthesize the first strand, we used converted DNA and 5′-biotinylated adaptor primers containing seven random nucleotides at its 3′ end (BioP5N7, biotin-ACACTCTTTCCCTACACGACGCTCTTCCGATCTNNNNNNN). The first strand product was purified using streptavidin-coated magnetic beads (Thermo Fisher Scientific, 11205D) and alkaline denaturation. Second strand DNA was synthesized using the immobilized first strand DNA and another adaptor primer also containing seven random nucleotides at its3′ end (P7N7, GTGACTGGAGTTCAGACGTGTGCTCTTCCGATCTNNNNNNN). Unique molecular barcodes and sequences essential for binding to Illumina flow-cells were added to the second strand DNA by PCR using 1 x HiFi HotStart Uracil+ Mix (KAPA, KK2801) and 10 µM indexed TruSeq-type oligos, amplified by 15 cycles of PCR and size selected by PEG-diluted SPRI beads. Library integrity was determined by agarose gel electrophoresis and sequenced on an Illumina HiSeq using single-end 100 bp chemistry.

**Bioinformatic analysis of the TET3-CD overexpression experiment**

The quality of the raw FASTQ files was evaluated using FastQC software [https://www.bioinformatics.babraham.ac.uk/projects/fastqc/] (v0.11.9). Raw reads were trimmed using Trim Galore! [https://www.bioinformatics.babraham.ac.uk/projects/trim_galore/] (v0.6.4), in a two-step process. First, adaptors were removed and 10 bp was hard-trimmed from the 5’ end of all reads and then low-quality base calls (Phred score < 20) were removed. Read mapping and base calling was performed using Bismark (v0.22.3) with the option --pbat specified (*11*). Mus musculus genome assembly GRCm38 (mm10) was used as reference. Bismark output files were deduplicated and methylation calls were obtained. The non-conversion rate during the bisulfite treatment was evaluated by calculating the proportion of non-CG methylation; by this measure, all libraries had a bisulfite conversion efficiency of at least 97.5% (Supplementary File 1).

Methylation in the CG context for each hexamer was calculated using an in-house Python script. Briefly, methylation calls were tracked to the reference genome, validated as CG context and the -2, -1, +1 and +2 nucleotides were examined to determine the identity of the CG-containing hexamer (NNCGNN). Then, methylation status for each call was extracted and added to a count table containing each hexamer. Finally, hexamer methylation was calculated as the proportion of total methylated cytosines over total cytosines. To examine methylation in CGI regions, previously published CGI coordinates (*12*) were converted from mm9 to GRCm38 (mm10) using the liftOver tool [https://genome.ucsc.edu/cgi-bin/hgLiftOver] (*12*). Then, methylation calls were classified as CGI or non-CGI and used as input for the methylation by hexamer script. Although some of CG-containing hexamers are more common than others in the mouse genome (meaning they were more likely to have overlapping reads associated with them), the average number of methylation calls for each motif was 6332. Theoretical asymptotic estimators predict technical error associated with this level of sequencing at ±1.03 p.p. (95% confidence) (*13*).

The demethylation velocity for each hexamer, following TET3-CD overexpression and decitabine treatment, was calculated using during the linear phase of demethylation (6-18 h and 0-32 h post-treatment, respectively). Further in-house R-scripts were developed to assess the characteristics of demethylation velocity for a range of motif parameters and groups (e.g., Intra-Motif Positional Preference and reverse complement motifs), all of which can be found on our GitHub code repository (https://github.com/TimHore-Otago/TET_specificity). Lastly, these demethylation velocities were compared to those calculated from previously published BS-seq datasets (Supplementary File 2), using the same workflow as outlined above.

### References


1. T. A. Hore *et al.*, *Proc. Natl. Acad. Sci. U.S.A.* 113, 12202–12207 (2016).
2. P. Oberacker *et al.*, *PLoS Biol.* 17, e3000107 (2019).
3. S. Kellner *et al.*, *Nucleic Acids Res*. 42, e142 (2014).
4. M. Krug, M. S. Weiss, U. Heinemann, U. Mueller, *J Appl Cryst*. 45, 568–572 (2012).
5. A. J. McCoy *et al.*, *J Appl Crystallogr*. 40, 658–674 (2007).
6. P. D. Adams *et al.*, *Methods*. 55, 94–106 (2011).

7. D. A. CASE *et al.*, *J Comput Chem*. 26, 1668–1688 (2005).

8. X. Chen *et al.*, *Genes Dis*. 2, 96–105 (2015).

9. X. Hu *et al.*, *Cell Stem Cell*. 14, 512–522 (2014).

10. J. R. Peat, O. Ortega-Recalde, O. Kardailsky, T. A. Hore, *F1000Res*. 6, 526 (2017).

11. F. Krueger, S. R. Andrews, *Bioinformatics*. 27, 1571–1572 (2011).

12. R. S. Illingworth *et al.*, *PLoS Genet.* 6, e1001134 (2010).

13. O. Ortega-Recalde *et al.*, *Annu Rev Anim Biosci*. 8, 47–69 (2020).

14. P. A. Ginno *et al.*, *Nat Commun*. 11, 2680 (2020).

15. J. Charlton *et al.*, *Nat. Genet.* 52, 819–827 (2020).

### Legends for Supplementary Figures

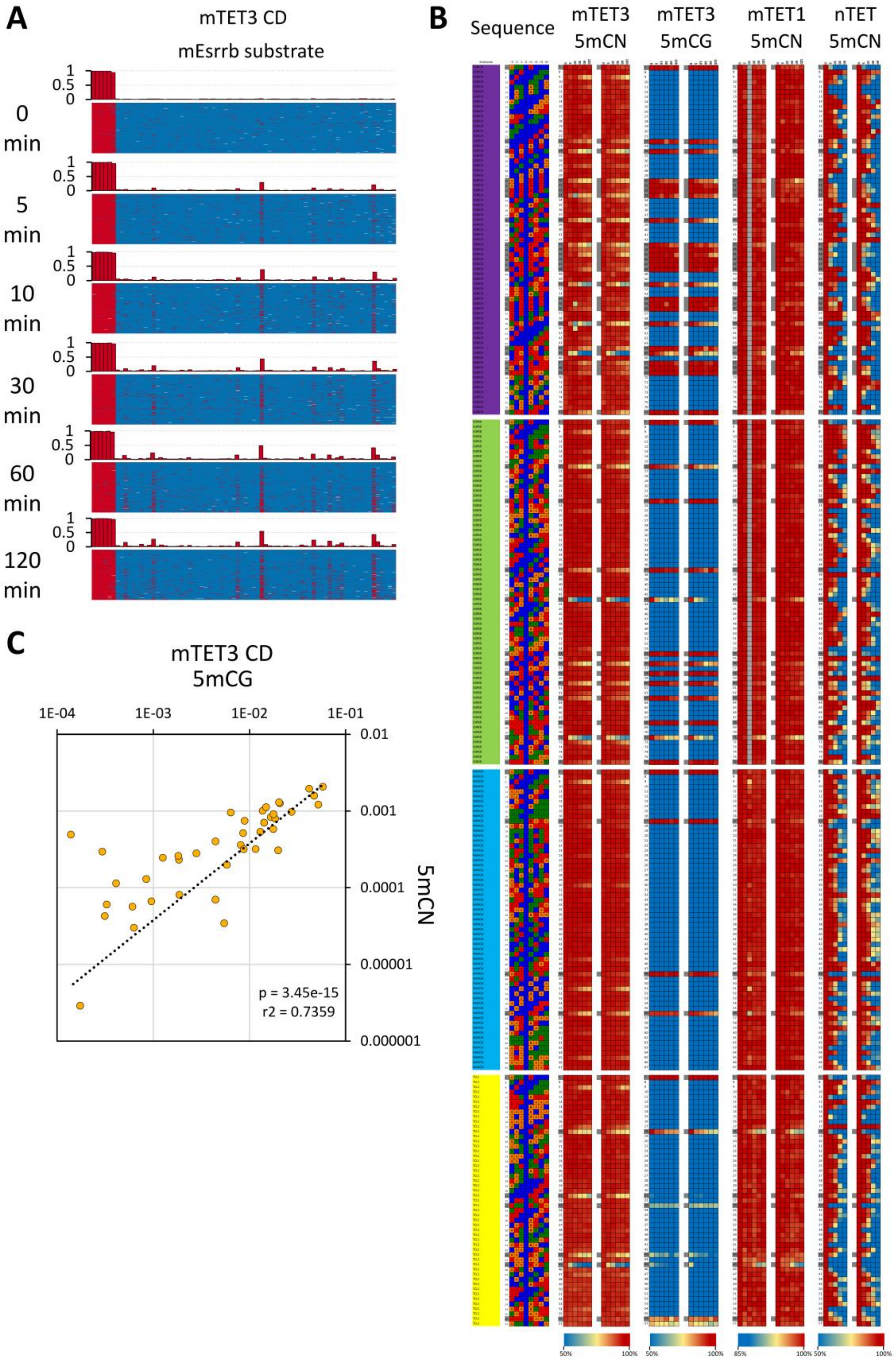

**Figure S1. Summary of the *in vitro* demethylation profiles. (A)** Example of primary results of single-molecule methylation profiles as obtained from the *in vitro* reactions for TET enzymes. The 5mC-modified substrates were incubated with mouse TET3 CD and the methylation profile was investigated on single molecules using bisulfite sequencing. Because each sequencing read stems from a single DNA molecule that is amplified in a cluster, the sequencing results illustrate the methylation state of that molecule. For each timepoint, the graphs represent 100 randomly selected sequences, rows denote separate DNA molecules, columns every possible C/5mC site present on the substrate. C/5fC/5caC are shown as red bars, 5mC/5hmC in blue. **(B)** Summary of the TET demethylation profiles obtained *in vitro*. Dots represent individual CG sites probed. Please note the different color scales for mTET1. The dsDNA substrate are indicated with colors: CGrich – violet, mEsrrb – green, mNanog – blue, mTcl1 – yellow. **(C)** Comparison between the mTET3 CD activity on the M.SssI methylated substrate (5mCG) and the 5mC modified substrate where all the Cs are replaced with 5mC (5mCN). X and Y axis represent fraction converted 5mC per minute.

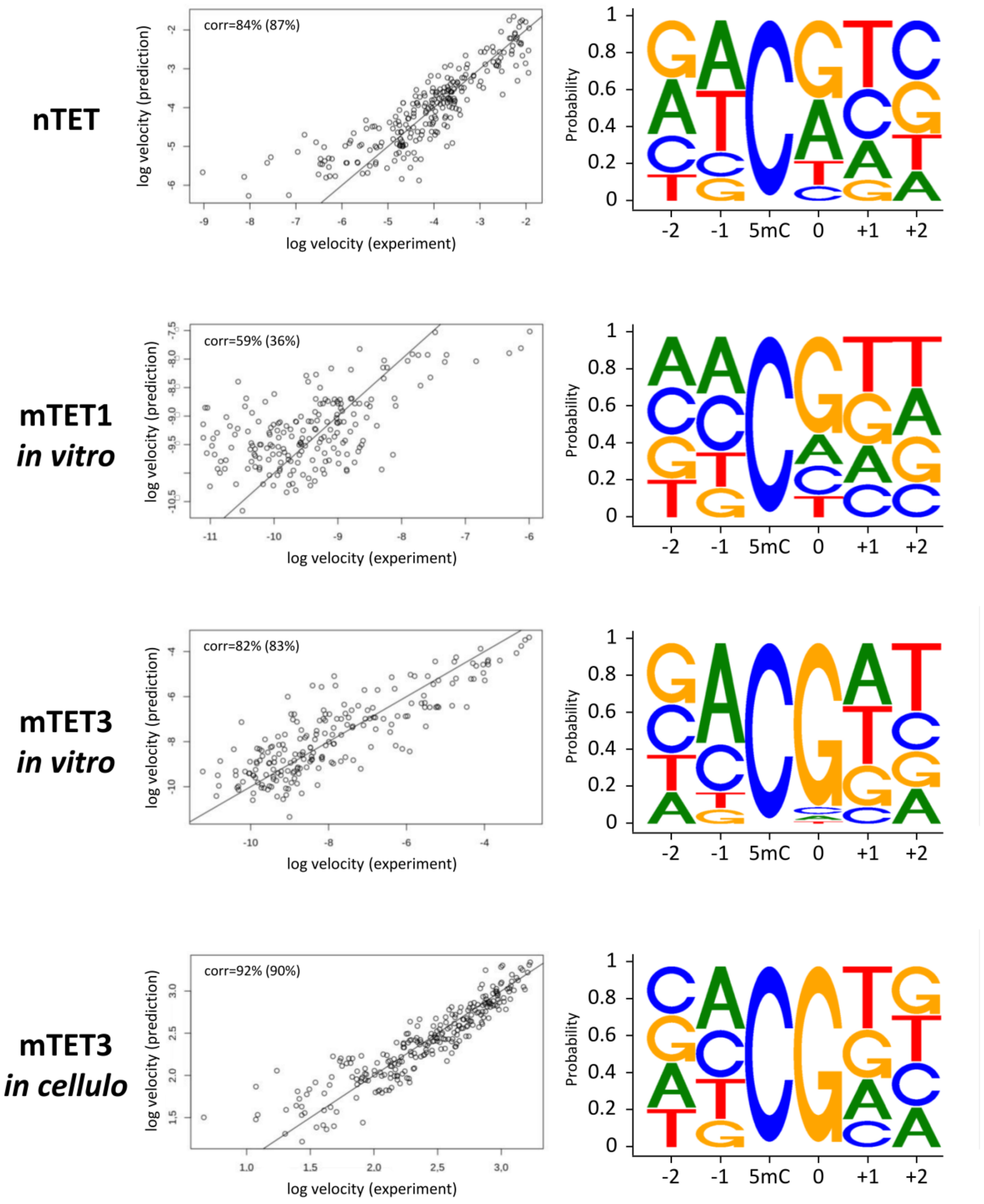


**Figure S2. Sequence models for the prediction of catalytic rates.** Reaction rates were analyzed in terms of a 5-base model (NNCNNN) for *in vitro* or a 4-base model (NNCGNN) for *in vivo* reaction velocities. The model assumes independent site preferences, equivalent to a linear model description for logarithmic reaction rates. Left panels show and quantify (R-values) the correlation between experimental and interpolated reaction velocities. R values are for a joint training and testing set, R values in brackets were calculated for data split in half into training and testing sets. The sequence logo representations are based on exponentials of regression coefficients. The logos in this figure represent all data, not just those for the most or least favorable substrates as in other figures and are therefore less pronounced. Noise in the weaker velocity data and the assumption of independent site preferences may also make the logos less clear-cut than those determined by other methods.

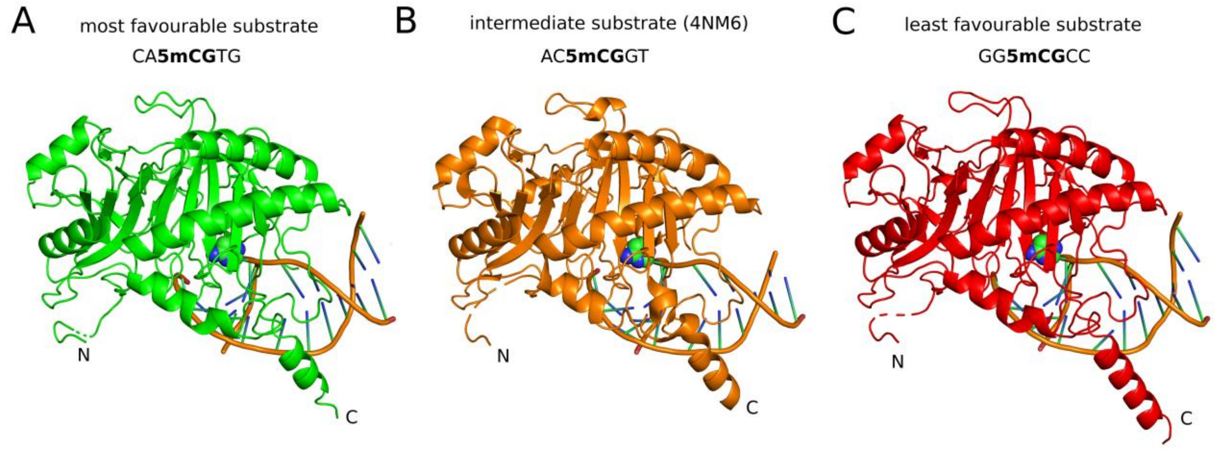


**Figure S3**. **Comparison of the hTET2 in complex with (A) the most favorable substrate, (B) an intermediate substrate, and (C) the least favorable substrate.** The apparent differences at the C-terminus (longer helix for the least optimal substrate) are not likely to be relevant. The B-factors are very high in this region of the protein, and the structures are actually similar (but happen to fall on either side of the boundaries for helix assignment). The genuine differences between the structures are not visible at this scale and are highlighted in Figure 3.

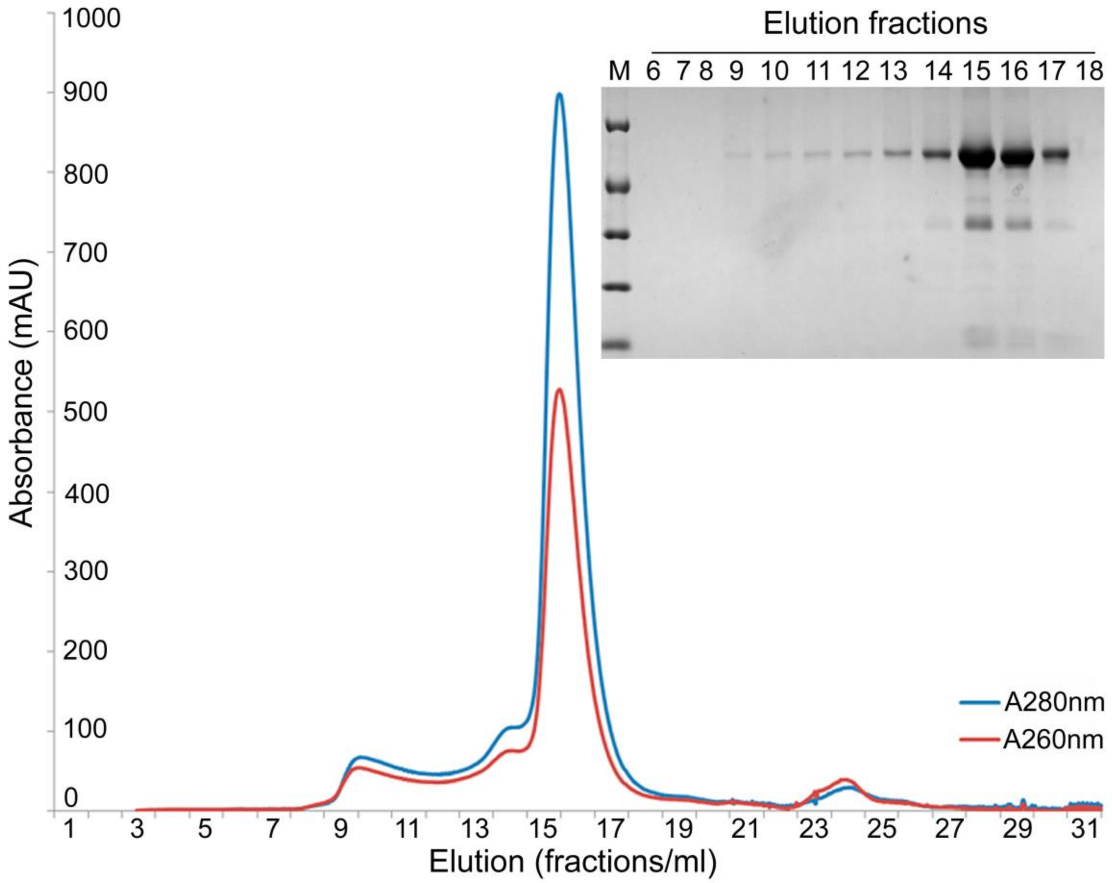


**Figure S4. Representative gel filtration profile of the hTET2 fragment used for crystallization.** The fragment has a calculated molecular mass of 51.4 kDa and unlike other TET preparations eluted in a single peak from a Superdex 200 column. Eluted fractions were analyzed by gel electrophoresis (12% PAGE) and stained by Coomassie Blue.

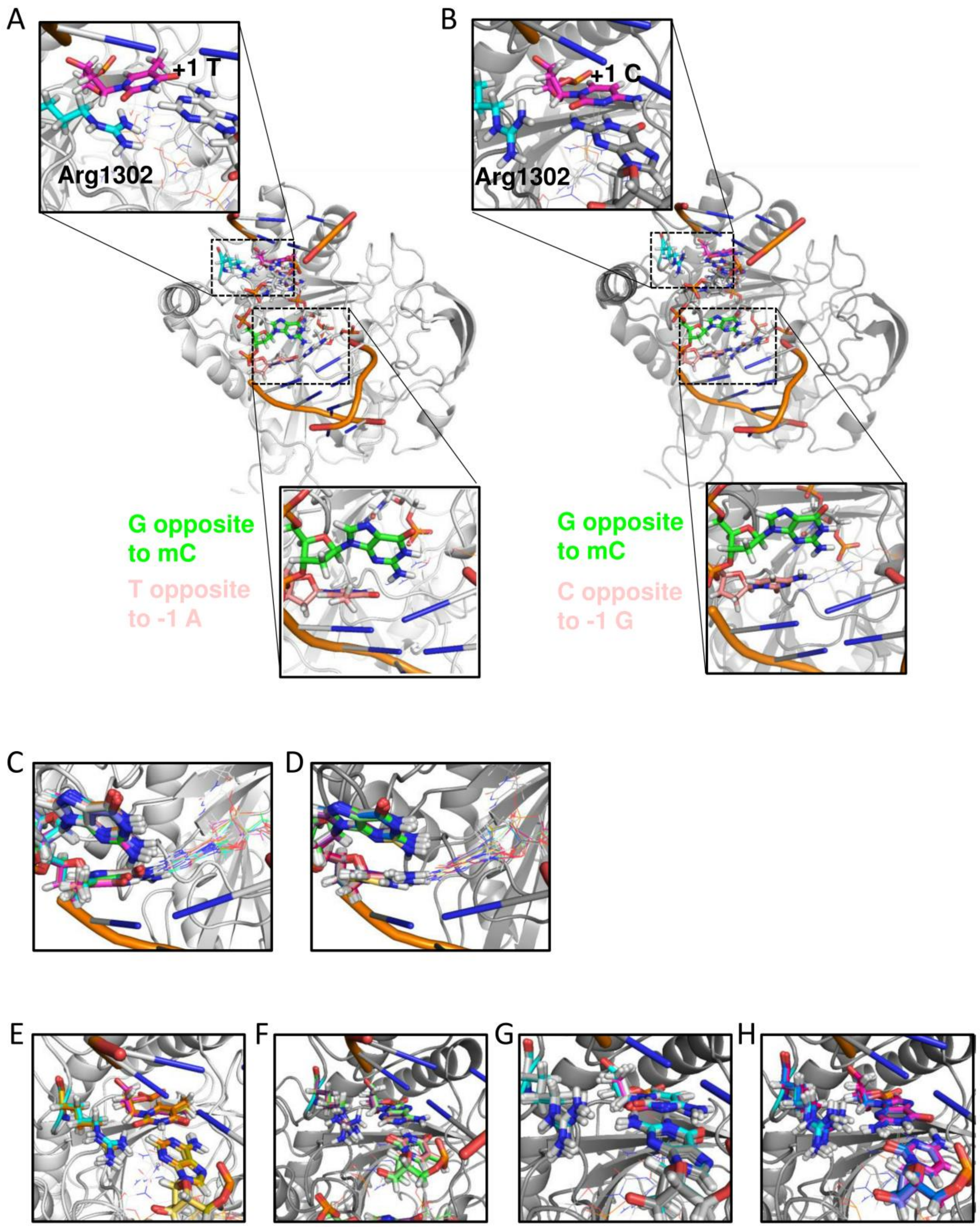


**Figure S5. Modeled complex between TET enzyme and target DNA. (A)** Complex with the most favorable substrate and **(B)** complex with the least favorable substrate. In both (A) and (B), insets on the top illustrate the interactions with Arg1302 that determine the 3'-preference. Insets at the bottom of the figure show the interactions in the non-substrate strand that determine the 5'-

preference. **(C,D)** Model explanation for the 5'-preference; **(C)** Superposition of all modeled DNA sequence variants in complex with TET with either a C or A at position -1 (in total 8 cases). For clarity only the guanine opposite to the flipped 5mC and the neighboring base (opposite to position -1) are shown as color-coded sticks **(D)** same as (C) but superposition of all modeled DNA sequence variants in complex with TET with either a T or G at position -1 (in total 8 cases). **(E-H)** Model explanation for the 3'-preference. Conformation and contacts of Arg1302 with the nucleotide at position 5 and the base on the opposite strand (stick model) **(E)** Superposition of all modeled DNA sequence variants in complex with TET with a T in position +1 (in total 4 cases). A stable hydrogen bond is formed. **(F)** same as (E) but with an A in position +1 and less ordered Arg1302 conformation. **(G)** same as (E) but with a C5, again no hydrogen bond contact with Arg1302 is formed, the side chain is dissociated from the DNA. **(H)** same as (G) but with a G in position +1.

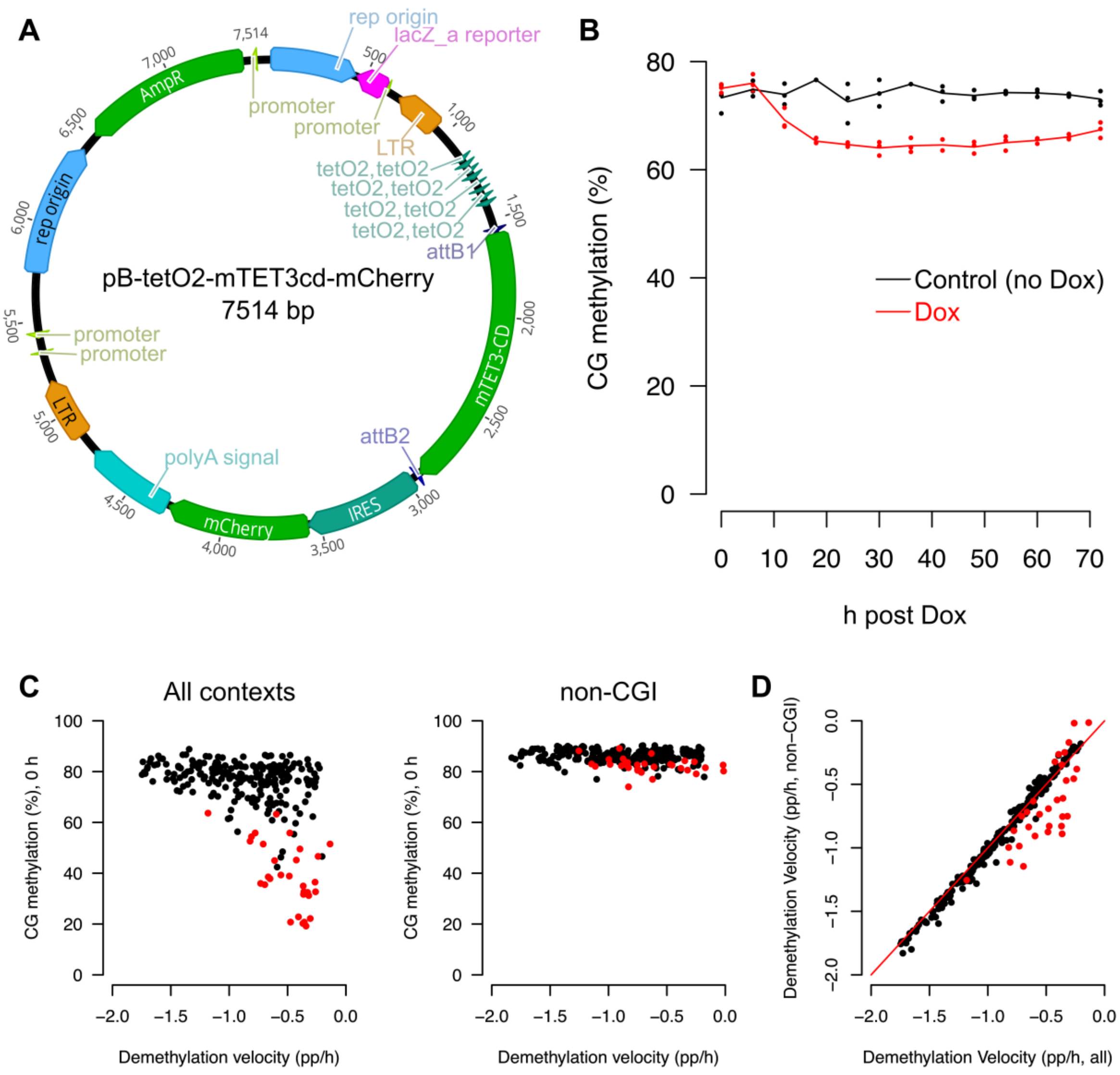


**Figure S6. TET3 catalytic domain overexpression in mouse embryonic stem cells.** In combination with helper plasmids (methods), the (**A**) pB-tetO2-meTET3cd-mCherry construct was used to drive (**B**) doxycycline-inducible loss of global CG methylation (Dox, red line), relative to controls (no Dox, black line). (**C**) Demethylation rate (percentage points per h, pp/h) of all 256 CG-containing hexamers, relative to their starting methylation levels in all contexts (left panel) and only non-CpG island (non-CGI) regions (right panel). (**D**) Demethylation rates of CG-containing hexamers in all contexts (x-axis) and the non-CGI contexts only (y-axis). For **C** and **D**, motifs with a CG in their sequence, in addition to the central CG, are highlighted in red.

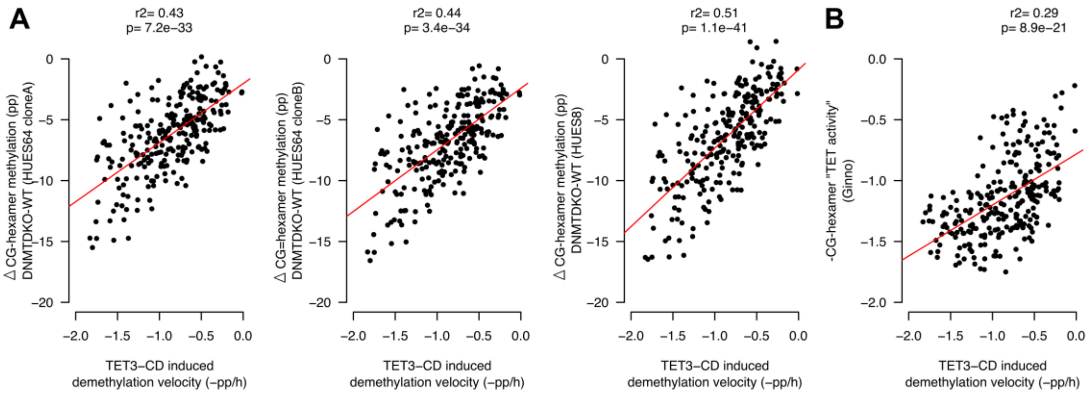


**Figure S7. TET hexamer preference can be detected in previously published data sets** (*14*)(*15*). Average demethylation velocity for all 256 CG-containing hexamers in mouse embryonic stem cells following overexpression of a TET3 catalytic domain (TET3-CD) transgene (x-axis, data from Figure 6) is significantly correlated with (**A**) the change in CG methylation loss at the same motifs genome-wide following knockout of DNMT3A and DNMT3B (DNMT-DKO) in 3 Human embryonic stem cell lines (HUES64-CloneA, HUES64-CloneB and HUES8) from Charlton et al., (2020). Also significantly correlated with TET3-CD induced demethylation velocity is (**B**) predicted TET-activity at 800,000 CG dinucleotides (Ginno et al., 2020) that were binned according to their hexamer identity.

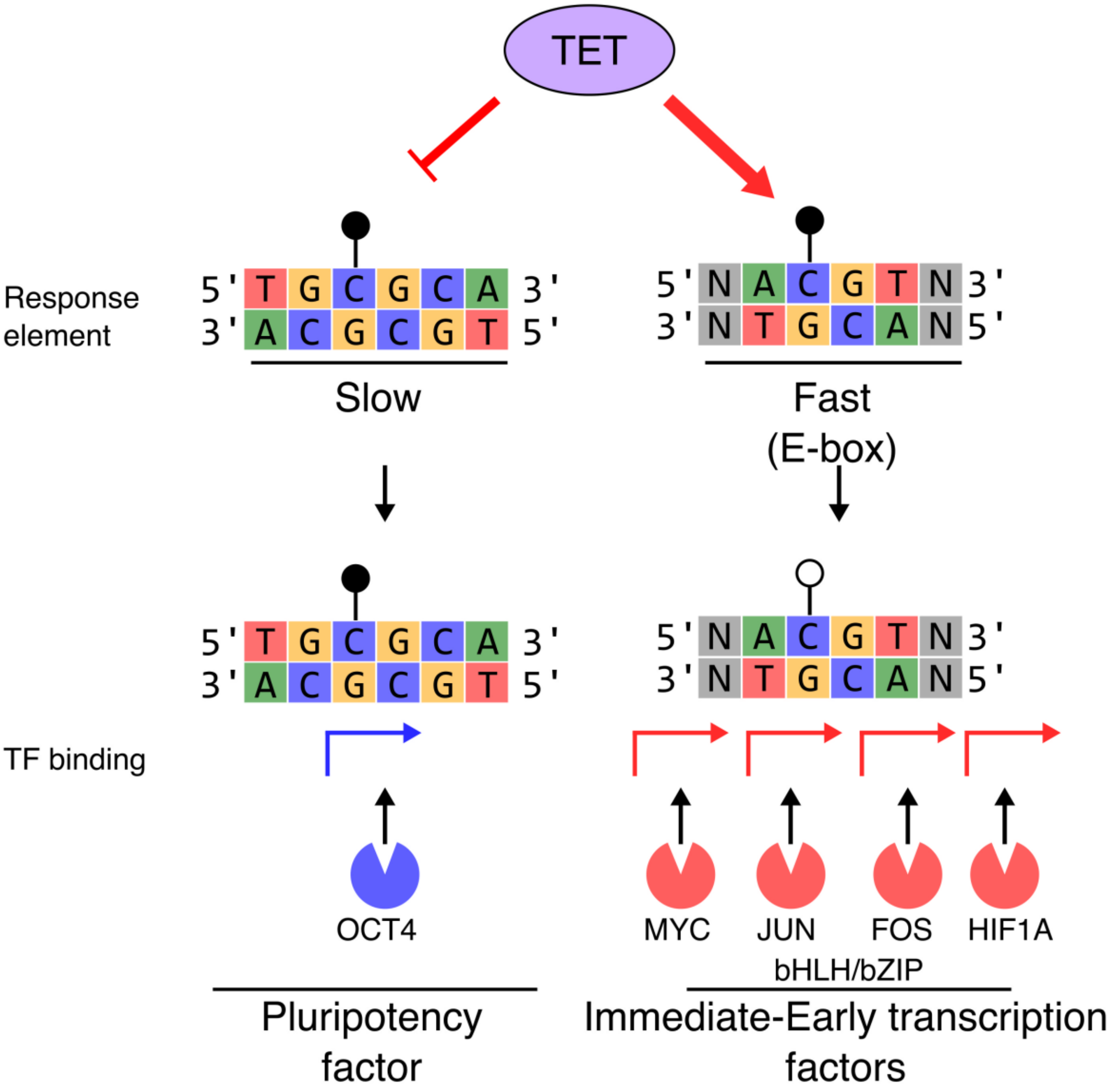


**Figure S8. Conceptual model.** The TET catalytic domain preferentially demethylates e-box sequences that bind immediately early transcription factors such as MYC, JUN, FOS and HIF1A. In contrast, TET does not rapidly demethylate other sequences such as TGCGCA, known to bind the pluripotency-related transcription factor OCT4 in the methylated state. Accordingly, TET sequence preference appears exquisitely tuned for its biological function supporting pluripotency and responding to environmental and biotic stimuli.

#### Tables S1 to S2

##### Table S1. Data collection and refinement statistics.

| | **TET2 favourable substrate 7NE3** | **TET2 unfavourable substrate 7NE6** |
|---|---|---|
| Resolution range (Å) | 43-2.26(2.3 - 2.26) | 43-2.3 (2.38 - 2.30) |
| Space group | $C222_1$ | $C222_1$ |
| Unit cell(Å), | 44.0,87.7,260.5 | 48.0, 88.0, 262.0 |
| Total reflections | 347,788 (32,571) | 189,003 (12,257) |
| Unique reflections | 26,450 (2,568) | 24,979 (2,337) |
| Multiplicity | 13.1 (12.7) | 7.6 (5.2) |
| Completeness (%) | 99.9 (99.2) | 99.2 (93.3) |
| I/σI | 16.4 (1.6) | 13.1 (1.74) |
| Wilson B-factor(Å$^2$) | 51.5 | 43.3 |
| $R_{merge}$(%) | 12.9 (157.1) | 11.7 (84.6) |
| $CC_{1/2}$ | 0.999 (0.689) | 0.997 (0.606) |
| $R_{work}$/$R_{free}$(%) | 17.5/22.6 | 19.5/22.7 |
| Number of non-hydrogen atoms | 3,969 | 4,040 |
| Macromolecules | 3,635 | 3,657 |
| Ligands | 74 | 70 |
| Solvent | 260 | 313 |
| Protein residues | 416 | 418 |
| RMSD bonds lengths (Å) | 0.008 | 0.003 |
| RMSD angles ($^o$) | 0.86 | 0.55 |
| Ramachandran favored (%) | 97.1 | 96.6 |
| Ramachandran allowed (%) | 2.7 | 3.2 |
| Ramachandran outliers (%) | 0.2 | 0.2 |
| Average B-factor(Å$^2$) | 63.0 | 51.6 |
| Macromolecules | 62.9 | 51.6 |
| Ligands | 67.0 | 51.7 |
| Solvent | 63.8 | 52.1 |

Statistics for the highest-resolution shell are shown in parentheses.

**Table S2. DNA fragments, oligonucleotides and primers used in this work.**

| Name | Sequence 5’ -> 3’ | |
|---|---|---|
| **Esrrb-F**<br>**Esrrb-R** | 5’-TGGTAGGGCTGCTGGACCTT-3’<br>5’-AAGCAATGCTGGTCATTCCCAC-3’ | Substrate amplification |
| **Nanog-F**<br>**Nanog-R** | 5’-AGCCACATTAGTTTATGTCTAAAAGTGTCT-3’<br>5’-GTGTGATATAATTAAACCAGCCATTTCCTTT-3’ | Substrate amplification |
| **Tcl1-F**<br>**Tcl1-R** | 5’-TAGGTCAGGGCCAAACACAAACAC-3’<br>5’-GCTACATCCATGTCTGCAGGG-3’ | Substrate amplification |
| **CGrich-F**<br>**CGrich-R** | 5’-CATCATCCCCAAGGCCTTCC-3’<br>5’-CCCTCCTCCTTCTCAATTTAACCC-3’ | Substrate amplification |
| **Synthetic substrates** | | |
| **Esrrb**<br>(GRCm38/mm10 chr12:86514434-8651468) | TGGTAGGGCTGCTGGACCTTTACCGAGCCATCCT<br>GCAGCTGGTGCGCAGGTACAAGAAACTCAAGGT<br>AGAGAAGGAAGAGTTTATGATCCTCAAGGCCCTG<br>GCCCTCGCCAACTCAGGTAAGGGTGGCACGTGAC<br>CCTCAGAGGGCTTCAGGGGTTCTGTGGGAGCCTA<br>ACAAGCGCCGCAAGCGCCGCACCGCTGCTCAGTC<br>TGTGGGCGGGGCTGGCTTACGGATGTGGGAATGA<br>CCAGCATTGCTT | *In vitro* dsDNA substrate<br>249 bp |
| **Nanog**<br>(GRCm38/mm10 chr6:122705362-122705621) | AGCCACATTAGTTTATGTCTAAAAGTGTCTAATT<br>GAAACAAGAAATGGCTGCTTTAGCCGGGTGTGGT<br>GGCACACGCCTTTAATCCCAGCACTCAGGAGGCA<br>GAGGCAGGCAGATTTCTGAGTTCAAGGCCAGCCT<br>GGTCTACAGAGTGAGTTCCAGGACAGCCAGAGCT<br>ACACAGAGAAACCCTGTCTCGAAAAACCAACCA<br>ACCAACCAACCAAACAAACAAACAAAAAAGGAA<br>ATGGCTGGTTTAATTATATCACAC | *In vitro* dsDNA substrate<br>260 bp |
| **Tcl1**<br>GRCm38/mm10 chr12:105217818-105218068) | TAGGTCAGGGCCAAACACAAACACCGTGAAGAG<br>AGTGGGGTGAAAAAAAAAAAAAGAGAGAGAAAT<br>AAAGAAAATATCCAATTTAAAAAAATCTCTGAGG<br>AAAAAAAAAAAAGAAGAAGAAATGGAGAAACCT<br>GCTTCAGTATTATCTGTGGATCCCCGGCTGAGTTC<br>GTGAGATGGCCCAGCAGTCATCTCTTAGACAACC<br>GATGGTGAACAACGAGTCACCTCAGGGGCCCTGC<br>AGACATGGATGTAGC | *In vitro* dsDNA substrate<br>251 bp |
| **CGrich_short**<br>(GRCh38/hg38 chr22: 22508772-22508926) | TAGTAACGGCCGCCAGTGTGCTGGAATTCGCCCT<br>TCATCATCCCCAAGGCCTTCCCCGCACGCCTCCA<br>CACGCGCGCGTCCAGTGGAGACCTGCGATTGGCT<br>GCCAGGTGCCGGCGCGAGATCGGCGCGGCTCCG<br>AGCTAGGAGCATGCGCGCGCTCTGACGCCCCTGG<br>TGGCGACGGCTGGACGCGGGGTTAAATTGAGAA<br>GGAGGAGGGAAGGGCGAATTCTGCAGATATCCA<br>TCACACTGGCGGCC | *In vitro* dsDNA substrate<br>249 bp |
| **LCMS_CACGTG** | 5’-GTAAGGGTGGCAmCGTGACCCTCAGAG-3’<br>5’-CTCTGAGGGTCACGTGCCACCCTTAC-3’ | LC/MS substrate<br>26 bp |
| **LCMS_GGCGGG** | 5’-GTAAGGGTGGGGmCGGGGCCCTCAGAG-3’<br>5’-CTCTGAGGGCCCCGCCCCACCCTTAC-3’ | LC/MS substrate<br>26 bp |
| **Kapa_f**<br>**Kapa_r** | 5’-AATGATACGGCGACCACCGA-3'<br>5’-CCGCAGAAGACGGCATACGA-3' | Library amplification |
